# Common variants in breast cancer risk loci predispose to distinct tumor subtypes

**DOI:** 10.1101/733402

**Authors:** Thomas U. Ahearn, Haoyu Zhang, Kyriaki Michailidou, Roger L. Milne, Manjeet K. Bolla, Joe Dennis, Alison M. Dunning, Michael Lush, Qin Wang, Irene L. Andrulis, Hoda Anton-Culver, Volker Arndt, Kristan J. Aronson, Paul L. Auer, Annelie Augustinsson, Adinda Baten, Heiko Becher, Sabine Behrens, Javier Benitez, Marina Bermisheva, Carl Blomqvist, Stig E. Bojesen, Bernardo Bonanni, Anne-Lise Børresen-Dale, Hiltrud Brauch, Hermann Brenner, Angela Brooks-Wilson, Thomas Brüning, Barbara Burwinkel, Saundra S. Buys, Federico Canzian, Jose E. Castelao, Jenny Chang-Claude, Stephen J. Chanock, Georgia Chenevix-Trench, Christine L. Clarke, NBCS Collaborators, J. Margriet Collée, Angela Cox, Simon S. Cross, Kamila Czene, Mary B. Daly, Peter Devilee, Thilo Dörk, Miriam Dwek, Diana M. Eccles, D. Gareth Evans, Peter A. Fasching, Jonine Figueroa, Giuseppe Floris, Manuela Gago-Dominguez, Susan M. Gapstur, José A. García-Sáenz, Mia M. Gaudet, Graham G. Giles, Mark S. Goldberg, Anna González-Neira, Grethe I. GrenakerAlnæs, Mervi Grip, Pascal Guénel, Christopher A. Haiman, Per Hall, Ute Hamann, Elaine F. Harkness, Bernadette A.M. Heemskerk-Gerritsen, Bernd Holleczek, Antoinette Hollestelle, Maartje J. Hooning, Robert N. Hoover, John L. Hopper, Anthony Howell, kConFab/AOCS Investigators, Milena Jakimovska, Anna Jakubowska, Esther M. John, Michael E. Jones, Audrey Jung, Rudolf Kaaks, Saila Kauppila, Renske Keeman, Elza Khusnutdinova, Cari M. Kitahara, Yon-Dschun Ko, Stella Koutros, Vessela N. Kristensen, Ute Krüger, Katerina Kubelka-Sabit, Allison W. Kurian, Kyriacos Kyriacou, Diether Lambrechts, Derrick G. Lee, Annika Lindblom, Martha Linet, Jolanta Lissowska, Ana Llaneza, Wing-Yee Lo, Robert J. MacInnis, Arto Mannermaa, Mehdi Manoochehri, Sara Margolin, Maria Elena Martinez, Catriona McLean, Alfons Meindl, Usha Menon, Heli Nevanlinna, William G. Newman, Jesse Nodora, Kenneth Offit, Håkan Olsson, Nick Orr, Tjoung-Won Park-Simon, Alpa V. Patel, Julian Peto, Guillermo Pita, Dijana Plaseska-Karanfilska, Ross Prentice, Kevin Punie, Katri Pylkäs, Paolo Radice, Gad Rennert, Atocha Romero, Thomas Rüdiger, Emmanouil Saloustros, Sarah Sampson, Dale P. Sandler, Elinor J. Sawyer, Rita K. Schmutzler, Minouk J. Schoemaker, Ben Schöttker, Mark E. Sherman, Xiao-Ou Shu, Snezhana Smichkoska, Melissa C. Southey, John J. Spinelli, Anthony J. Swerdlow, Rulla M. Tamimi, William J. Tapper, Jack A. Taylor, Lauren R. Teras, Mary Beth Terry, Diana Torres, Melissa A. Troester, Celine M. Vachon, Carolien H.M. van Deurzen, Elke M. van Veen, Philippe Wagner, Clarice R. Weinberg, Camilla Wendt, Jelle Wesseling, Robert Winqvist, Alicja Wolk, Xiaohong R. Yang, Wei Zheng, Fergus J. Couch, Jacques Simard, Peter Kraft, Douglas F. Easton, Paul D.P. Pharoah, Marjanka K. Schmidt, Montserrat García-Closas, Nilanjan Chatterjee

**Affiliations:** Division of Cancer Epidemiology and Genetics, National Cancer Institute, National Institutes of Health, Department of Health and Human Services, Bethesda, MD, USA; Department of Biostatistics, Johns Hopkins Bloomberg School of Public Health, Baltimore, MD, USA; Biostatistics Unit, The Cyprus Institute of Neurology & Genetics, Nicosia, Cyprus; Centre for Cancer Genetic Epidemiology, Department of Public Health and Primary Care, University of Cambridge, Cambridge, UK; Cyprus School of Molecular Medicine, The Cyprus Institute of Neurology & Genetics, Nicosia, Cyprus; Cancer Epidemiology Division, Cancer Council Victoria, Melbourne, Victoria, Australia; Centre for Epidemiology and Biostatistics, Melbourne School of Population and Global Health, The University of Melbourne, Melbourne, Victoria, Australia; Precision Medicine, School of Clinical Sciences at Monash Health, Monash University, Clayton, Victoria, Australia; Centre for Cancer Genetic Epidemiology, Department of Oncology, University of Cambridge, Cambridge, UK; Fred A. Litwin Center for Cancer Genetics, Lunenfeld-Tanenbaum Research Institute of Mount Sinai Hospital, Toronto, ON, Canada; Department of Molecular Genetics, University of Toronto, Toronto, ON, Canada; Department of Medicine, Genetic Epidemiology Research Institute, University of California Irvine, Irvine, CA, USA; Division of Clinical Epidemiology and Aging Research, German Cancer Research Center (DKFZ), Heidelberg, Germany; Department of Public Health Sciences, and Cancer Research Institute, Queen’s University, Kingston, ON, Canada; Cancer Prevention Program, Fred Hutchinson Cancer Research Center, Seattle, WA, USA; Zilber School of Public Health, University of Wisconsin-Milwaukee, Milwaukee, WI, USA; Department of Cancer Epidemiology, Clinical Sciences, Lund University, Lund, Sweden; Leuven Multidisciplinary Breast Center, Department of Oncology, Leuven Cancer Institute, University Hospitals Leuven, Leuven, Belgium; Institute of Medical Biometry and Epidemiology, University Medical Center Hamburg-Eppendorf, Hamburg, Germany; Division of Cancer Epidemiology, German Cancer Research Center (DKFZ), Heidelberg, Germany; Human Cancer Genetics Programme, Spanish National Cancer Research Centre (CNIO), Madrid, Spain; Biomedical Network on Rare Diseases (CIBERER), Madrid, Spain; Institute of Biochemistry and Genetics, Ufa Federal Research Centre of the Russian Academy of Sciences, Ufa, Russia; Saint Petersburg State University, Saint-Petersburg, Russia; Department of Oncology, Helsinki University Hospital, University of Helsinki, Helsinki, Finland; Department of Oncology, Örebro University Hospital, Örebro, Sweden; Faculty of Health and Medical Sciences, University of Copenhagen, Copenhagen, Denmark; Department of Clinical Biochemistry, Herlev and Gentofte Hospital, Copenhagen University Hospital, Herlev, Denmark; Copenhagen General Population Study, Herlev and Gentofte Hospital, Copenhagen University Hospital, Herlev, Denmark; Division of Cancer Prevention and Genetics, IEO, European Institute of Oncology IRCCS, Milan, Italy; Department of Cancer Genetics, Institute for Cancer Research, Oslo University Hospital-Radiumhospitalet, Oslo, Norway; Institute of Clinical Medicine, Faculty of Medicine, University of Oslo, Oslo, Norway; Dr. Margarete Fischer-Bosch-Institute of Clinical Pharmacology, Stuttgart, Germany; iFIT-Cluster of Excellence, University of Tübingen, Tübingen, Germany; German Cancer Consortium (DKTK) and German Cancer Research Center (DKFZ), Partner Site Tübingen, Tübingen, Germany; German Cancer Consortium (DKTK), German Cancer Research Center (DKFZ), Heidelberg, Germany; Division of Preventive Oncology, German Cancer Research Center (DKFZ) and National Center for Tumor Diseases (NCT), Heidelberg, Germany; Genome Sciences Centre, BC Cancer Agency, Vancouver, BC, Canada; Department of Biomedical Physiology and Kinesiology, Simon Fraser University, Burnaby, BC, Canada; Institute for Prevention and Occupational Medicine of the German Social Accident Insurance, Institute of the Ruhr University Bochum (IPA), Bochum, Germany; Molecular Epidemiology Group, C080, German Cancer Research Center (DKFZ), Heidelberg, Germany; Molecular Biology of Breast Cancer, University Womens Clinic Heidelberg, University of Heidelberg, Heidelberg, Germany; Department of Medicine, Huntsman Cancer Institute, Salt Lake City, UT, USA; Genomic Epidemiology Group, German Cancer Research Center (DKFZ), Heidelberg, Germany; Oncology and Genetics Unit, Instituto de Investigacion Sanitaria Galicia Sur (IISGS), Xerencia de Xestion Integrada de Vigo-SERGAS, Vigo, Spain; Cancer Epidemiology Group, University Cancer Center Hamburg (UCCH), University Medical Center Hamburg-Eppendorf, Hamburg, Germany; Department of Genetics and Computational Biology, QIMR Berghofer Medical Research Institute, Brisbane, Queensland, Australia; Westmead Institute for Medical Research, University of Sydney, Sydney, New South Wales, Australia; Department of Clinical Genetics, Erasmus University Medical Center, Rotterdam, The Netherlands; Sheffield Institute for Nucleic Acids (SInFoNiA), Department of Oncology and Metabolism, University of Sheffield, Sheffield, UK; Academic Unit of Pathology, Department of Neuroscience, University of Sheffield, Sheffield, UK; Department of Medical Epidemiology and Biostatistics, Karolinska Institutet, Stockholm, Sweden; Department of Clinical Genetics, Fox Chase Cancer Center, Philadelphia, PA, USA; Department of Human Genetics, Leiden University Medical Center, Leiden, The Netherlands; Department of Pathology, Leiden University Medical Center, Leiden, The Netherlands; Gynaecology Research Unit, Hannover Medical School, Hannover, Germany; School of Life Sciences, University of Westminster, London, UK; Faculty of Medicine, University of Southampton, Southampton, UK; North West Genomics Laboratory Hub, Manchester Centre for Genomic Medicine, St Mary’s Hospital, Manchester University NHS Foundation Trust, Manchester Academic Health Science Centre, Manchester, UK; Division of Evolution and Genomic Sciences, School of Biological Sciences, Faculty of Biology, Medicine and Health, University of Manchester, Manchester Academic Health Science Centre, Manchester, UK; David Geffen School of Medicine, Department of Medicine Division of Hematology and Oncology, University of California at Los Angeles, Los Angeles, CA, USA; Usher Institute of Population Health Sciences and Informatics, The University of Edinburgh, Edinburgh, UK; Cancer Research UK Edinburgh Centre, The University of Edinburgh, Edinburgh, UK; Fundación Pública Galega de Medicina Xenómica, Instituto de Investigación Sanitaria de Santiago de Compostela (IDIS), Complejo Hospitalario Universitario de Santiago, SERGAS, Santiago de Compostela, Spain; Moores Cancer Center, University of California San Diego, La Jolla, CA, USA; Behavioral and Epidemiology Research Group, American Cancer Society, Atlanta, GA, USA; Medical Oncology Department, Hospital Clínico San Carlos, Instituto de Investigación Sanitaria San Carlos (IdISSC), Centro Investigación Biomédica en Red de Cáncer (CIBERONC), Madrid, Spain; Division of Clinical Epidemiology, Royal Victoria Hospital, McGill University, Montréal, QC, Canada; Department of Medicine, McGill University, Montréal, QC, Canada; Department of Surgery, Oulu University Hospital, University of Oulu, Oulu, Finland; Center for Research in Epidemiology and Population Health (CESP), Team Exposome and Heredity, INSERM, University Paris-Saclay, Villejuif, France; Department of Preventive Medicine, Keck School of Medicine, University of Southern California, Los Angeles, CA, USA; Department of Oncology, Södersjukhuset, Stockholm, Sweden; Molecular Genetics of Breast Cancer, German Cancer Research Center (DKFZ), Heidelberg, Germany; Division of Informatics, Imaging and Data Sciences, Faculty of Biology, Medicine and Health, University of Manchester, Manchester Academic Health Science Centre, Manchester, UK; Nightingale & Genesis Prevention Centre, Wythenshawe Hospital, Manchester University NHS Foundation Trust, Manchester, UK; NIHR Manchester Biomedical Research Unit, Manchester University NHS Foundation Trust, Manchester Academic Health Science Centre, Manchester, UK; Department of Medical Oncology, Erasmus MC Cancer Institute, Rotterdam, The Netherlands; Saarland Cancer Registry, Saarbrücken, Germany; Division of Cancer Sciences, University of Manchester, Manchester, UK; Research Centre for Genetic Engineering and Biotechnology “Georgi D. Efremov”, MASA, Skopje, Republic of North Macedonia; Department of Genetics and Pathology, Pomeranian Medical University, Szczecin, Poland; Independent Laboratory of Molecular Biology and Genetic Diagnostics, Pomeranian Medical University, Szczecin, Poland; Department of Epidemiology & Population Health, Stanford University School of Medicine, Stanford, CA, USA; Department of Medicine, Division of Oncology, Stanford Cancer Institute, Stanford University School of Medicine, Stanford, CA, USA; Division of Genetics and Epidemiology, The Institute of Cancer Research, London, UK; Department of Pathology, Oulu University Hospital, University of Oulu, Oulu, Finland; Division of Molecular Pathology, The Netherlands Cancer Institute - Antoni van Leeuwenhoek Hospital, Amsterdam, The Netherlands; Department of Genetics and Fundamental Medicine, Bashkir State University, Ufa, Russia; Radiation Epidemiology Branch, Division of Cancer Epidemiology and Genetics, National Cancer Institute, Bethesda, MD, USA; Department of Internal Medicine, Evangelische Kliniken Bonn gGmbH, Johanniter Krankenhaus, Bonn, Germany; Department of Medical Genetics, Oslo University Hospital and University of Oslo, Oslo, Norway; Department of Histopathology and Cytology, Clinical Hospital Acibadem Sistina, Skopje, Republic of North Macedonia; Department of Electron Microscopy/Molecular Pathology, The Cyprus Institute of Neurology & Genetics, Nicosia, Cyprus; Laboratory for Translational Genetics, Department of Human Genetics, University of Leuven, Leuven, Belgium; VIB Center for Cancer Biology, Leuven, Belgium; Cancer Control Research, BC Cancer, Vancouver, BC, Canada; Department of Mathematics and Statistics, St. Francis Xavier University, Antigonish, NS, Canada; Department of Molecular Medicine and Surgery, Karolinska Institutet, Stockholm, Sweden; Department of Clinical Genetics, Karolinska University Hospital, Stockholm, Sweden; Department of Cancer Epidemiology and Prevention, M. Sklodowska-Curie Cancer Center, Oncology Institute, Warsaw, Poland; General and Gastroenterology Surgery Service, Hospital Universitario Central de Asturias, Oviedo, Spain; University of Tübingen, Tübingen, Germany; Institute of Clinical Medicine, Pathology and Forensic Medicine, University of Eastern Finland, Kuopio, Finland; Translational Cancer Research Area, University of Eastern Finland, Kuopio, Finland; Biobank of Eastern Finland, Kuopio University Hospital, Kuopio, Finland; Department of Clinical Science and Education, Södersjukhuset, Karolinska Institutet, Stockholm, Sweden; Anatomical Pathology, The Alfred Hospital, Melbourne, Victoria, Australia; Department of Gynecology and Obstetrics, University of Munich, Campus Großhadern, Munich, Germany; Institute of Clinical Trials & Methodology, University College London, London, UK; Department of Obstetrics and Gynecology, Helsinki University Hospital, University of Helsinki, Helsinki, Finland; Herbert Wertheim School of Public Health and Human Longevity Science, University of California San Diego, La Jolla, CA, USA; Clinical Genetics Research Lab, Department of Cancer Biology and Genetics, Memorial Sloan Kettering Cancer Center, New York, NY, USA; Centre for Cancer Research and Cell Biology, Queen’s University Belfast, Belfast, Ireland, UK; Department of Non-Communicable Disease Epidemiology, London School of Hygiene and Tropical Medicine, London, UK; Human Genotyping-CEGEN Unit, Human Cancer Genetic Program, Spanish National Cancer Research Centre, Madrid, Spain; Department of General Medical Oncology and Multidisciplinary Breast Center, Leuven Cancer Institute, University Hospitals Leuven, Leuven, Belgium; Laboratory of Cancer Genetics and Tumor Biology, Cancer and Translational Medicine Research Unit, Biocenter Oulu, University of Oulu, Oulu, Finland; Laboratory of Cancer Genetics and Tumor Biology, Northern Finland Laboratory Centre Oulu, Oulu, Finland; Unit of Molecular Bases of Genetic Risk and Genetic Testing, Department of Research, Fondazione IRCCS Istituto Nazionale dei Tumori (INT), Milan, Italy; Clalit National Cancer Control Center, Carmel Medical Center and Technion Faculty of Medicine, Haifa, Israel; Medical Oncology Department, Hospital Universitario Puerta de Hierro, Madrid, Spain; Institute of Pathology, Staedtisches Klinikum Karlsruhe, Karlsruhe, Germany; Department of Oncology, University Hospital of Larissa, Larissa, Greece; Prevent Breast Cancer Centre and Nightingale Breast Screening Centre, Manchester University NHS Foundation Trust, Manchester, UK; Epidemiology Branch, National Institute of Environmental Health Sciences, NIH, Research Triangle Park, NC, USA; School of Cancer & Pharmaceutical Sciences, Comprehensive Cancer Centre, Guy’s Campus, King’s College London, London, UK; Center for Integrated Oncology (CIO), Faculty of Medicine and University Hospital Cologne, University of Cologne, Cologne, Germany; Center for Molecular Medicine Cologne (CMMC), Faculty of Medicine and University Hospital Cologne, University of Cologne, Cologne, Germany; Center for Familial Breast and Ovarian Cancer, Faculty of Medicine and University Hospital Cologne, University of Cologne, Cologne, Germany; Network Aging Research, University of Heidelberg, Heidelberg, Germany; Department of Health Sciences Research, Mayo Clinic College of Medicine, Jacksonville, FL, USA; Division of Epidemiology, Department of Medicine, Vanderbilt Epidemiology Center, Vanderbilt-Ingram Cancer Center, Vanderbilt University School of Medicine, Nashville, TN, USA; Ss. Cyril and Methodius University in Skopje, Medical Faculty, University Clinic of Radiotherapy and Oncology, Skopje, Republic of North Macedonia; Department of Clinical Pathology, The University of Melbourne, Melbourne, Victoria, Australia; Population Oncology, BC Cancer, Vancouver, BC, Canada; School of Population and Public Health, University of British Columbia, Vancouver, BC, Canada; Division of Breast Cancer Research, The Institute of Cancer Research, London, UK; Department of Population Health Sciences, Weill Cornell Medicine, New York, NY, USA; Epigenetic and Stem Cell Biology Laboratory, National Institute of Environmental Health Sciences, NIH, Research Triangle Park, NC, USA; Department of Epidemiology, Mailman School of Public Health, Columbia University, New York, NY, USA; Institute of Human Genetics, Pontificia Universidad Javeriana, Bogota, Colombia; Department of Epidemiology, Gillings School of Global Public Health and UNC Lineberger Comprehensive Cancer Center, University of North Carolina at Chapel Hill, Chapel Hill, NC, USA; Department of Health Science Research, Division of Epidemiology, Mayo Clinic, Rochester, MN, USA; Department of Pathology, Erasmus University Medical Center, Rotterdam, The Netherlands; Biostatistics and Computational Biology Branch, National Institute of Environmental Health Sciences, NIH, Research Triangle Park, NC, USA; Department of Pathology, The Netherlands Cancer Institute - Antoni van Leeuwenhoek hospital, Amsterdam, The Netherlands; Institute of Environmental Medicine, Karolinska Institutet, Stockholm, Sweden; Department of Surgical Sciences, Uppsala University, Uppsala, Sweden; Department of Laboratory Medicine and Pathology, Mayo Clinic, Rochester, MN, USA; Genomics Center, Centre Hospitalier Universitaire de Québec, Université Laval Research Center, Department of Molecular Medicine, Université Laval, Québec City, QC, Canada; Program in Genetic Epidemiology and Statistical Genetics, Harvard T.H. Chan School of Public Health, Boston, MA, USA; Department of Epidemiology, Harvard T.H. Chan School of Public Health, Boston, MA, USA; Division of Psychosocial Research and Epidemiology, The Netherlands Cancer Institute - Antoni van Leeuwenhoek hospital, Amsterdam, The Netherlands; Department of Biostatistics, Bloomberg School of Public Health, John Hopkins University, Baltimore, MD, USA; Department of Oncology, School of Medicine, John Hopkins University, Baltimore, MD, USA

**Keywords:** Breast Cancer, etiologic heterogeneity, genetic predisposition, common breast cancer susceptibility variants

## Abstract

**Background:** Genome-wide association studies (GWAS) have identified multiple common breast cancer susceptibility variants. Many of these variants have differential associations by estrogen receptor (ER), but how these variants relate with other tumor features and intrinsic molecular subtypes is unclear.

**Methods:** Among 106,571 invasive breast cancer cases and 95,762 controls of European ancestry with data on 173 breast cancer variants identified in previous GWAS, we used novel two-stage polytomous logistic regression models to evaluate variants in relation to multiple tumor features (ER, progesterone receptor (PR), human epidermal growth factor receptor 2 (HER2) and grade) adjusting for each other, and to intrinsic-like subtypes.

**Results:** Eighty-five of 173 variants were associated with at least one tumor feature (false discovery rate <5%), most commonly ER and grade, followed by PR and HER2. Models for intrinsic-like subtypes found nearly all of these variants (83 of 85) associated at P<0.05 with risk for at least one luminal-like subtype, and approximately half (41 of 85) of the variants were associated with risk of at least one non-luminal subtype, including 32 variants associated with triple-negative (TN) disease. Ten variants were associated with risk of all subtypes in different magnitude. Five variants were associated with risk of luminal A-like and TN subtypes in opposite directions.

**Conclusion:** This report demonstrates a high level of complexity in the etiology heterogeneity of breast cancer susceptibility variants and can inform investigations of subtype-specific risk prediction.

## Introduction

Breast cancer represents a heterogenous group of diseases with different molecular and clinical features[1]. Clinical assessment of estrogen receptor (ER), progesterone receptor (PR), human epidermal growth factor receptor 2 (HER2) and histological grade are routinely determined to inform treatment strategies and prognostication[2]. Combined, these tumor features define five intrinsic-like subtypes (i.e., luminal A-like, luminal B–like/HER2-negative, luminal B-like/HER2-positive, HER2-positive/non-luminal, and triple negative) that are correlated with intrinsic subtypes defined by gene expression panels[2, 3]. Most known breast cancer risk or protective factors are related to luminal or hormone receptor (ER or PR) positive tumors, whereas less is known about the etiology of triple-negative (TN) tumors, an aggressive subtype[4, 5].

Breast cancer genome-wide association studies (GWAS) have identified over 170 common susceptibility variants, most of them single nucleotide polymorphisms (SNPs), of which many are differentially associated with ER-positive than ER-negative disease[6–8]. These include 20 variants that primarily predispose to ER-negative or TN disease[7, 8]. However, few studies have evaluated variant associations with other tumor features, or simultaneously studied multiple, correlated tumor markers to identify source(s) of etiologic heterogeneity[7, 9–13]. We recently developed a two-stage polytomous logistic regression method that efficiently characterizes etiologic heterogeneity while accounting for tumor marker correlations and missing tumor data[14, 15]. This method can help describe complex relationships between susceptibility variants and multiple tumor features, helping to clarify breast cancer subtype etiologies and increasing the power to generate more accurate risk estimates between susceptibility variants and less common subtypes. We recently demonstrated the power of this method in a GWAS to identify novel breast cancer susceptibility accounting for tumor heterogeneity[15].

In this report, we sought to expand our understanding of etiologic heterogeneity among breast cancer subtypes, by applying the two-stage polytomous logistic regression methodology to a large study population from the Breast Cancer Association Consortium (BCAC) for detailed characterization of risk associations with 173 breast cancer risk variants identified by GWAS[6, 7] by tumor subtypes defined by ER, PR, HER2 and tumor grade.

## Methods

### Study Population and Genotyping

The study population and genotyping are described in previous publications[6, 7] and in the **Supplementary Methods**. We included invasive cases and controls from 81 BCAC studies with genotyping data from two Illumina genome-wide custom arrays, the iCOGS and OncoArray (106,571 cases (OncoArray: 71,788; iCOGS: 34,783) and 95,762 controls (OncoArray: 58,134; iCOGS: 37,628); **Supplementary Table 1**). We evaluated 173 breast cancer risk variants that were identified in or replicated by prior BCAC analyses to be associated with breast cancer risk at a p-value threshold p<5.0×10^-8^ [6, 7]. Most of these variants (n=153) were identified because of their association with risk of overall breast cancer, and a small number of variants (n=20) were identified because of their association specific to ER-negative breast cancer (**Supplementary Table 2**). These 173 variants have not previously been simultaneously investigated for evidence of tumor heterogeneity with multiple tumor markers[6, 7, 15, 16]. Genotypes for the variants marking the 173 susceptibility loci were determined by genotyping with the iCOGS and the OncoArray arrays and imputation to the 1000 Genomes Project (Phase 3) reference panel.

### Statistical Analysis

An overview of the analytic strategy is shown in **Figure 1** and a detailed discussion of the statistical methods, including the two-stage polytomous logistic regression, are provided in the **Supplementary Methods** and elsewhere[14, 15]. Briefly, we used two-stage polytomous regression models that allow modelling of genetic association of breast cancer accounting for underlying heterogeneity in associations by combinations of multiple tumor markers using a parsimonious decomposition of subtype-specific case-control odds-ratio parameters in terms of marker-specific case-case odd-ratio parameters[14, 15]. We introduced further parsimony by using mixed-effect formulation of the model that allows ER-specific case-case parameters to be treated as fixed and similar parameters for other markers (PR, HER2 and grade) as random. We used an expectation–maximization (EM) algorithm[17] for parameter estimation under this model to account for missing data in tumor characteristics. A detailed description of the two-stage polytomous regression models used in this manuscript is presented in the **Supplementary Methods** and in separate manuscripts[14, 15].

**Figure 1.**
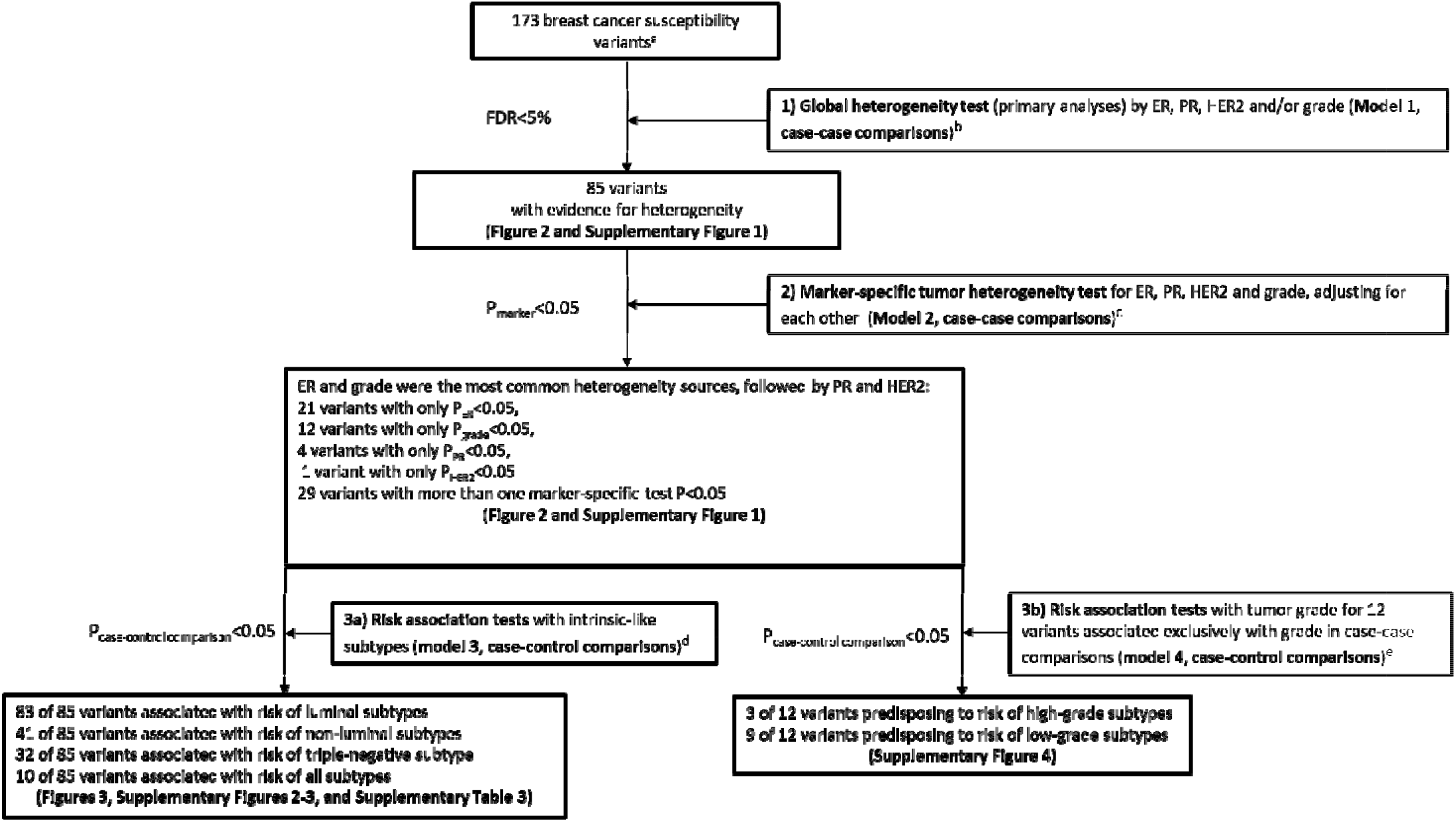
Overview of the analytic strategy and results from the investigation of 173 known breast cancer susceptibility variants for evidence of heterogeneity according to the estrogen receptor (ER), progesterone receptor (PR), human epidermal growth factor receptor 2 (HER2), and grade ^a^We evaluated 173 breast cancer risk variants identified in or replicated by prior BCAC GWAS [6, 7], see **Methods** and **Supplementary Methods** sections for more details. ^b^ Model 1 (primary analyses): Mixed-effect two-stage polytomous model (ER as fixed-effect, and PR, HER2 and grade as random-effects) for <u>global heterogeneity tests</u> (i.e. case-case comparisons from stage 2 of the two-stage model) between each individual risk variant and any of the tumor features (separate models were fit for each variant). ^c^ Model 2: Fixed-effect two-stage polytomous model for <u>marker-specific tumor heterogeneity tests</u> (i.e. case-case comparisons from stage 2 of the two-stage model) between each individual variant and each of the tumor features (ER, PR, HER2, and grade), mutually adjusted for each other (separate models were fit for each variant). ^d^ Model 3: Fixed effect two-stage polytomous model <u>for risk associations with intrinsic-like subtypes</u> (i.e. case-control comparisons from stage 1 of the two-stage model): luminal A-like, luminal B-like/HER2-negative, luminal B-like/HER2-positive, HER2-positive/non-luminal, and triple negative. ^e^ Model 4: Fixed effect two-stage polytomous model <u>for risk associations with tumor grade</u> (i.e. case-control comparisons from stage 1 of the two-stage model) for the 12 variants associated at P<0.05 only with grade in case-case comparisons (from model 2): grade 1, grade 2, and grade 3.

Our primary aim was to identify which of 173 known breast cancer susceptibility variants showed heterogenous risk associations by ER, PR, HER2 and grade. This was tested using a global heterogeneity test by ER, PR, HER2 and/or grade, with a mixed-effect two-stage polytomous model (**model 1**), fitted separately for each variant. The global null hypothesis was that there was no difference in risk of breast cancer associated with the variant genotype across any of the tumor features being evaluated. We accounted for multiple testing (173 tests, one for each of variant) of the global heterogeneity test using a false discovery rate (FDR) <0.05% under the Benjamini-Hochberg procedure[18].

For the variants showing evidence of global heterogeneity after FDR adjustment, we further evaluated which of the tumor features contributed to the heterogeneity by fitting a fixed-effects two-stage model (**model 2**) that simultaneously tested for associations with each tumor feature (this model was fitted for each variant separately). We used a threshold of P<0.05 for marker-specific tumor heterogeneity tests to describe which specific tumor marker(s) contributed to the observed heterogeneity, adjusting for the other tumor markers in the model. This p-value threshold was used only for descriptive purposes, as the primary hypotheses were tested using the FDR-adjusted global test for heterogeneity described above.

We conducted additional analyses to explore evidence of heterogeneity. We fitted a fixed-effect two-stage model (**model 3**) to estimate case-control odd ratios (ORs) and 95% confidence intervals (CI) between the variants and five intrinsic-like subtypes defined by combinations of ER, PR, HER2 and grade: (1) luminal A-like (ER+ and/or PR+, HER2-, grade 1 or 2); (2) luminal B-like/HER2-negative (ER+ and/or PR+, HER2-, grade 3); (3) luminal B-like/HER2-positive (ER+ and/or PR+, HER2+); (4) HER2-positive/non-luminal (ER- and PR-, HER2+), and (5) TN (ER-, PR-, HER2-). We also fitted a fixed-effect two-stage model to estimate case-control ORs and 95% confidence intervals (CI) with tumor grade (**model 4**; defined as grade 1, grade 2, and grade 3) for the variants associated at P<0.05 only with grade in case-case comparisons from model 2.

To help describe sources of heterogeneity from different tumor characteristics in models 2 and 3, we performed cluster analyses based on Euclidean distance calculated from z-statistics that were estimated by the individual marker-specific tumor heterogeneity tests (model 2) and the case-control associations with risk of intrinsic-like subtypes (model 3). The clusters were used only to help present our findings and were not intended to suggest strictly defined categories. Clustering was performed in R using the function Heatmap as implemented by the package “Complex Heatmap” version 3.1[19].

We performed sensitivity analyses, in which we estimated the ORs and 95% CI between the variants and the intrinsic-like subtypes by implementing a standard polytomous model restricted to cases with complete tumor marker data. For all analyses we analyzed OncoArray and iCOGS array data separately, adjusting for the first 10 principal components for ancestry-informative variants, and then meta-analyzed the results.

## Results

The mean (SD) ages at diagnosis (cases) and enrollment (controls) were 56.6 (12.2) and 56.4 (12.2) years, respectively. Among cases with information on the corresponding tumor marker, 81% were ER-positive, 68% PR-positive, 83% HER2-negative and 69% grade 1 or 2 (**Table 1;** see **Supplementary Table 1** for details by study). **Supplementary Table 3** shows the correlation between the tumor markers. ER was positively correlated with PR (r^2^=0.61) and inversely correlated with HER2 (r^2^=-0.16) and grade (r^2^=-0.39). The most common intrinsic-like subtype was luminal A-like (54%), followed by TN (14%), luminal B-like/HER2-negative (13%), Luminal B-like/HER2-positive (13%) and HER2-positive/non-luminal (6%; **Table 1**).

**Table 1.** Distribution of estrogen receptor (ER), progesterone receptor (PR), human epidermal growth factor receptor 2 (HER2), and grade and the intrinsic-like subtypes^a^ among cases of invasive breast cancer in studies from the Breast Cancer Consortium Association.

| Tumor Marker |  | N (%) |
| --- | --- | --- |
| ER | Negative | 16,900 (19%) |
|  | Positive | 70,030 (81%) |
|  | Unknown | 19,641 |
| PR | Negative | 24,283 (32%) |
|  | Positive | 51,603 (68%) |
|  | Unknown | 30,685 |
| HER2 | Negative | 47,693 (83%) |
|  | Positive | 9,529 (17%) |
|  | Unknown | 49,349 |
| Grade | 1 | 15,583 (20%) |
|  | 2 | 37,568 (49%) |
|  | 3 | 24,382 (31%) |
|  | Unknown | 29,038 |
| Intrinsic-like subtypes |  |  |
| Luminal A-like |  | 27,510 (54%) |
| Luminal B-like/HER2-negative |  | 6,804 (13%) |
| Luminal B-like/HER2-positive |  | 6,511 (13%) |
| HER2-positive/non-luminal |  | 2,797 (6%) |
| Triple-negative |  | 7,178 (14%) |
| Unknown |  | 55,771 |

a Luminal A-like (ER+ and/or PR+, HER2-, grade 1 & 2); Luminal B-like/HER2-negative (ER+ and/or PR+, HER2-, grade 3); Luminal B-like/HER2-positive (ER+ and/or PR+, HER2+); HER2-positive/non-luminal (ER- and PR-, HER2+), and triple-negative (ER-, PR-, HER2-)

**Figure 1** shows an overview of the analytic strategy and results from three main analyses performed separately for each variant: 1) global test for heterogeneity by all tumor markers (model 1; primary hypothesis), 2) marker-specific tumor test for heterogeneity for each marker, adjusting for the others (model 2), and 3) estimation of case-control ORs (95%CIs) by intrinsic-like subtypes (model 3).

### 1) Global test for heterogeneity by tumor markers (primary hypothesis)

Mixed-effects two-stage models (model 1) were fitted for each of the 173 variants separately and included terms for ER, PR, HER2 and grade to test of global heterogeneity by any of the tumor features (case-case comparison). This model identified 85 of 173 (49.1%) variants with evidence of heterogeneity by at least one tumor feature (FDR<5%; **Figure 1–2; Supplementary Figure 1**). **Figure 2** shows a heatmap of the second stage z-values from the fixed-effects two-stage polytomous model for the marker-specific heterogeneity tests (case-case comparison from model 2) for the 173 breast cancer susceptibility variants and ER, PR, HER2, and grade.

**Figure 2.**
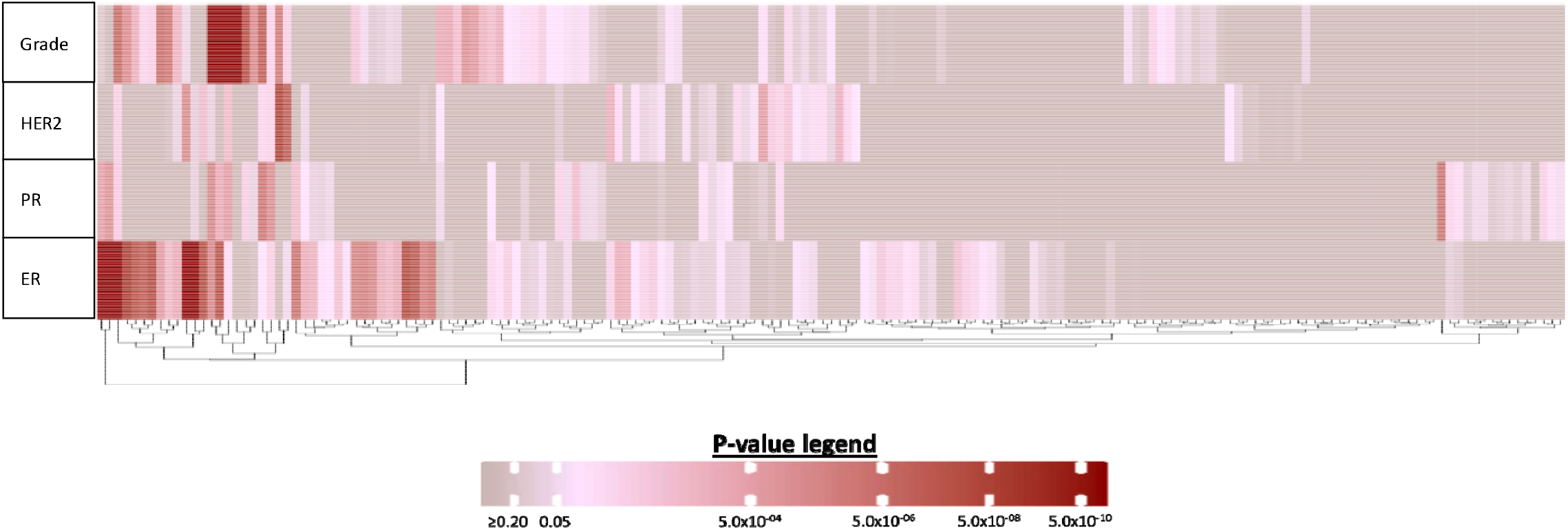
Heatmap of the z-values from the fixed-effects two-stage polytomous model for marker-specific heterogeneity tests (case-case comparison from model 2) for association between each of the 173 breast cancer susceptibility variants and estrogen receptor (ER), progesterone receptor (PR), human epidermal growth factor receptor 2 (HER2) or grade, adjusting for principal components and each tumor marker. Columns represent individual variants. For more detailed information on the context of figure see **Supplementary Figure 1**.

### 2) Marker-specific tumor test for heterogeneity for each marker, adjusting for other markers

Fixed-effects two-stage models (model 2) were used to test which of the correlated tumor markers was responsible for the observed global heterogeneity in the 85 variants (case-case comparison). These analyses identified ER and grade as the two features that most often contributed to the observed heterogeneity (45 and 33 variants had marker-specific P<0.05 for ER and grade, respectively), and 29 variants were associated with more than one tumor feature (**Figure 1–2**, **Supplementary Figure 1**). Eighteen of these 85 variants showed no associations with any individual tumor marker at P<0.05 (**Supplementary Figure 1**). Twenty-one variants were associated at P<0.05 only with ER, 12 variants only with grade, 4 variants only with PR and one variant only with HER2 (**Supplementary Figure 1,** see footnotes).

### 3) Estimation of case-control ORs (95%CIs) by intrinsic-like subtypes (model 3)

Fixed-effects two-stage models for intrinsic-like subtypes (model 3) were fitted for each of the 85 variants with evidence of global heterogeneity to estimate ORs (95% CIs) for risk associations with each subtype (case-control comparisons). **Supplementary Figure 2** shows a summary of these analyses for the 85 variants, clustered by case-control z-value of association between susceptibility variants and breast cancer intrinsic-like subtypes, and **Supplementary Figure 3** shows forest plots for associations with risk by tumor subtypes. Nearly all (83 of 85) variants were associated with risk (P<0.05) for at least one luminal-like subtype, and approximately half (41 of 85) of the variants were associated with risk of at least one nonluminal subtype, including 32 variants that were associated with TN disease (**Figure 1**, **Supplementary Figure 2 footnote ‘h’**). Ten variants were associated with risk of all subtypes (**Figure 1**, **Supplementary Figure 2 footnote ‘j’**). Below we describe examples of groups of variants associated with different patterns of associations with intrinsic subtypes (**Figure 3 a-d**).

**Figure 3.**
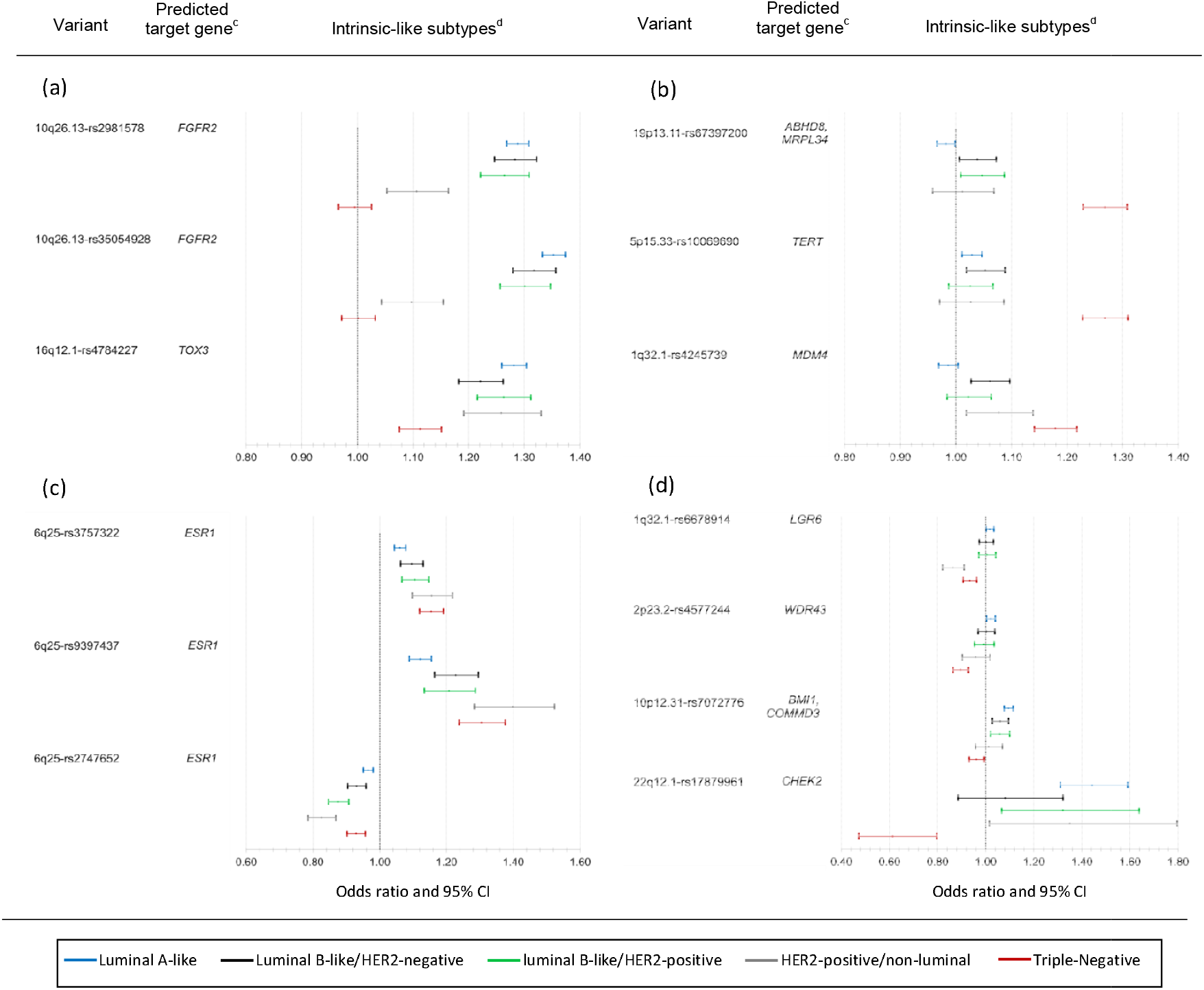
Results from fixed-effects two-stage polytomous models for risk associations^a^ with intrinsic-like subtypes (model 3) for variants with evidence of heterogeneity by tumor markers in the two-stage model (model 1)^b^; panels show examples of variants (a) most strongly associated with luminal-like subtypes, (b) most strongly associated with TN subtypes, (c) associated with all subtypes with varying strengths of association, and (d) associated with luminal A-like and TN subtypes in different directions. See **Supplementary Figure 3** for more details. ^a^ Per-minor allele odds ratio (95% confidence limits). ^b^ Model 1, mixed-effects two-stage polytomous model testing for global heterogeneity according to estrogen receptor (ER), progesterone receptor (PR), human epidermal growth factor receptor 2 (HER2) and grade ^c^ Predicted target genes as reported in Fachal L, et al. Nature genetics 2020; 52 (1), 56-73 ^d^ Luminal A-like (ER+ and/or PR+, HER2-, grade 1 & 2); Luminal B-like/HER2-negative (ER+ and/or PR+, HER2-, grade 3); luminal B-like/HER2-positive (ER+ and/or PR+, HER2+); HER2-positive/non-luminal (ER- and PR-, HER2+), and triple-negative (ER-, PR-, HER2-)

Two correlated (r^2^=0.73) variants at 10q26.13 (rs2981578 and rs35054928) and 16q12.1-rs4784227 had the strongest evidence of association with risk of luminal-like subtypes (**Figure 3a**, **Supplementary Figure 2**). The two variants at 10q26.13 showed no evidence of associations with TN subtypes, and a weaker association with HER2-positive/non-luminal subtype (**Figure 3a**, **Supplementary Figure 2**). In contrast, 16q12.1-rs4784227 was strongly associated with risk for all luminal-like subtypes and, weaker so, with risk of HER2-positive/non-luminal and TN subtypes (**Figures 3a**, **Supplementary Figure 2**).

Three variants 19p13.11-rs67397200, 5p15.33-rs10069690 and 1q32.11-rs4245739 showed the strongest evidence of associations with risk of TN disease (**Figure 3b**, **Supplementary Figure 2**).

Two weakly correlated variants in 6q25 (r^2^=0.17), rs9397437 and rs3757322, and a third variant in 6q25, rs2747652, which was not correlated (r^2^<0.01) with rs9397437 or rs3757322, showed strong evidence of being associated with risk of all subtypes, rs9397437 and rs3757322 had strong evidence of associations with risk of TN and risk of luminal-like subtypes (**Figures 3c** and **Supplementary Figure 2**). rs2747652 was most strongly associated with risk of HER2-positive subtypes (**Figures 3c**, **Supplementary Figure 2**).

Five variants were associated with risk of luminal A-like disease in an opposite direction to their association with risk of TN disease (**Figure 3d**, **Supplementary Figure 2**). 1q32.1-rs6678914, 2p23.2-rs4577244, and 19p13.11-rs67397200 had weaker evidence of associations with risk of luminal A-like disease compared to associations with risk of TN disease, and 10p12.31-rs7072776 and 22q12.1-rs17879961 (I157T) had stronger evidence of an association with risk of luminal A-like disease compared to their association with risk of TN disease (**Figure 3d**, **Supplementary Figure 2**, for rs67397200 see **Figure 3b**).

### 4) Estimation of case-control ORs (95%CIs) by tumor grade (model 4)

Case-control associations by tumor grade for the 12 variants associated at P<0.05 only with grade in case-case comparisons are shown in **Supplementary Figure 4**. 13q13.1-rs11571833, 1p22.3-rs17426269 and 11q24.3-rs11820646 showed stronger evidence for predisposing to risk of high-grade subtypes, and the remaining variants showed stronger evidence for predisposing to risk of low-grade subtypes.

When limiting analyses to cases with complete tumor marker data, results from casecontrol analyses were similar, but less precise than results from the two-stage polytomous regression model using the EM algorithm to account for missing tumor marker data (**Supplementary Table 4**).

## Discussion

This study demonstrates the extent and complexity of genetic etiologic heterogeneity among 173 breast cancer risk variants by multiple tumor characteristics, using novel methodology in the largest and the most comprehensive investigation conducted to date. We found compelling evidence that about half of the investigated breast cancer susceptibility loci (85 of 173 variants) predispose to tumors with different characteristics. We identified tumor grade, along with confirming ER status, as important determinants of etiologic heterogeneity. Associations with individual tumor features translated into differential associations with the risk of intrinsic-like subtypes defined by their combinations.

Many of the variants with evidence of global heterogeneity predisposed to risk of multiple subtypes, but with different magnitudes. For example, three variants identified in early GWAS for overall breast cancer, *FGFR2* (rs35054928 and rs2981578)[20, 21] and 8q24.21 (rs13281615)[20], were associated with luminal-like and HER2-positive/non-luminal subtypes, but not with TN disease. rs4784227 located near *TOX3[20, 22]* and rs62355902 located in a *MAP3K1[20]* regulatory element, were associated with risk of all five subtypes. Of the five variants found associated in opposite directions with luminal A-like and TN disease, we previously reported rs6678914 and rs4577244 to have opposite effects between ER-negative and ER-positive tumors[7]. rs17879961 (I157T), a likely causal[16] mis-sense variant located in a *CHEK2* functional domain that reduces or abolishes substrate binding[23], was previously reported to have opposite directions of effects on lung adenocarcinoma and lung squamous cell carcinoma and for lung cancer between smokers and non-smokers[24, 25]. However, further studies of these five variants are required to follow-up and clarify the mechanisms for these apparent cross-over effects.

In prior ER-negative GWAS, we identified 20 variants that predispose to ER-negative disease, of which five variants were only or most strongly associated with risk of TN disease (rs4245739, rs10069690, rs74911261, rs11374964, and rs67397200)[7, 8]. We confirmed these five variants to be most strongly associated with TN disease. The remaining previously identified 15 variants all showed associations with risk of non-luminal subtypes, especially TN disease, and for all but four variants (rs17350191, rs200648189, rs6569648, and rs322144) evidence of global heterogeneity was observed.

Little is known regarding PR and HER2 as sources of etiologic heterogeneity independent of ER status. Of the four variants that showed evidence of heterogeneity only according to PR, rs10759243[6, 26], rs11199914[27] and rs72749841[6] were previously found primarily associated with risk of ER-positive disease, and rs10816625 was found to be associated with risk of ER-positive/PR-positive tumors, but not other ER/PR combinations[12]. rs10995201 was the only variant found in case-case comparisons to be solely associated with HER2 status, although the evidence was not strong, requiring further confirmation. Previously, rs10995201 showed no evidence of being associated with ER status[28]. Most variants associated with PR or HER2, had not been investigated for PR or HER2 heterogeneity while adjusting for ER[9–13]. We previously reported rs10941679 to be associated with PR-status, independent of ER, and also with grade[10]. We also found suggestive evidence of PR-specific heterogeneity for 16q12-rs3803662[13], which is in high LD (r^2^= 0.78) with rs4784227 (*TOX3*), a variant strongly associated with PR status. Our findings for rs2747652 are also consistent with a prior BCAC fine-mapping analysis across the *ESR1* locus, which found rs2747652 to be associated with risk of the HER2-positive/non-luminal subtype and high grade independent of ER[9]. rs2747652 overlaps an enhancer region and is associated with reduced *ESR1* and *CCDC170* expression[9].

Histologic grade is a composite of multiple tumor characteristics including mitotic count, nuclear pleomorphism, and degree of tubule or gland formation[29]. Among the 12 variants identified with evidence of heterogeneity by grade only, rs17426269, rs11820646, and rs11571833 were found to be most strongly associated with risk of grade 3 disease. rs11571833 lies in the *BRCA2* coding region and produces a truncated form of the protein[30] and has been shown to be associated with both risk of TN disease and risk of serous ovarian tumors, both of which tend to be high-grade[31]. To our knowledge, rs17426269 and rs11820646 have not been investigated in relation to grade heterogeneity. The remaining 9 variants were all more strongly associated with grade 1 or grade 2 disease. Six of these variants were previously reported to be associated primarily with ER-positive disease[6, 27, 32, 33], highlighting the importance of accounting for multiple tumor characteristics to better illuminate heterogeneity sources.

We identified 18 variants with evidence of global heterogeneity (FDR<5%), but no significant (marker-specific P<0.05) associations with any of the individual tumor characteristic(s). This is likely explained by the fact that the test for association with specific tumor markers using fixed-effects models are less powerful than mixed-effects models used to test the primary hypothesis of global heterogeneity by any tumor marker[14].

To help describe and visualize the strength of the evidence for common heterogeneity patterns, we performed clustered analyses of z-values for tumor marker-specific heterogeneity tests and case-control associations with risk of intrinsic-like subtypes. Because they are based on z-values, these clusters reflect differences in sample size and statistical power to detect associations between variants and specific tumor subtypes. Thus, clusters should not be interpreted as strictly defined categories.

A major strength of our study is our large sample size of over 100,000 breast cancer cases with tumor marker information, and a similar number of controls, making this the largest, most comprehensive breast cancer heterogeneity investigation. Our application of the two-stage polytomous logistic regression enabled adjusting for multiple, correlated tumor markers and accounting for missing tumor marker data. This is a more powerful and efficient modeling strategy for identifying heterogeneity sources among highly correlated tumor markers, compared with standard polytomous logistic regression[14, 15]. In simulated and real data analyses, we have demonstrated that in the presence of heterogenous associations across subtypes, the two-stage model is more powerful than polytomous logistic regression for detecting heterogeneity. Moreover, we have demonstrated that in the presence of correlated markers, the two-stage model, incorporating all markers simultaneously, has much better ability to distinguish the true source(s) of heterogeneity compared to testing for heterogeneity by analysis of one marker at a time[14, 15]. In prior analyses, we showed that the two-stage polytomous regression is a powerful approach to identify susceptibility variants that display tumor heterogeneity[15]. Notably, in this prior investigation we excluded the genomic regions in which the 173 variants that were investigated in this work are located[15].

Our study also has some limitations. First, many breast cancer cases from studies included in this report had missing information on one or more tumor characteristics. ER tumor status data was available for 81% of cases, but missing data for the other tumor markers ranged from 27% to 46%. To address this limitation, we implemented an EM algorithm that allowed a powerful analysis to incorporate cases with missing tumor characteristics under the assumption that tumor characteristics are *missing at random* (MAR), i.e., the underlying reason for missing data may depend on observed tumor markers or/and covariate values, but not on the missing values themselves[34]. If this assumption is violated it can lead to an inflated type-one error[14]. However, in the context of genetic association testing, the missingness mechanism would also need to be related to the genetic variants under study, which is unlikely. Our study focused on investigating ER, PR, HER2, and grade as heterogeneity sources, and future studies with more detailed tumor characterization could reveal additional etiologic heterogeneity sources.

In summary, our findings provide insights into the complex etiologic heterogeneity patterns of common breast cancer susceptibility loci. These findings may inform future studies, such as fine-mapping and functional analyses to identify the underlying causal variants, clarifying biological mechanisms that drive genetic predisposition to breast cancer subtypes. Moreover, these analyses provide precise estimates of relative risk for different intrinsic-like subtypes that could improve the discriminatory accuracy of subtype-specific polygenic risk scores [35].

## Supporting information

Fundings and acknowledgement

## Abbreviations

GWAS: Genome-wide association studies
ER: estrogen receptor
PR: progesterone receptor
HER2: human epidermal growth factor receptor 2
FDR: false discovery rate
TN: triple-negative
BCAC: Breast Cancer Association Consortium
EM: expectation–maximization
OR: odd ratios
95% CI: 95% confidence interval

## Declarations

### Ethics approval and consent to participate

All the studies included in these analyses were approved by local IRBs.

### Consent for publication

Not applicable

### Availability of data and materials

Requests for data can be made to M.K.B. (; http://bcac.ccge.medschl.cam.ac.uk/bcacdata/).

### Author Contributions

Writing group: T.U.A., H.Z., K.Mi., R.L.M., F.J.C., J.Si., P.Kr., D.F.E., P.D.P.P., M.K.S., M.G-C., N.Ch.; Statistical analysis: H.Z., T.U.A., M.G-C., N.Ch.; Provision of DNA samples and/or phenotypic data: K.Mi., R.L.M., M.K.B., J.Den., A.M.D., M.Lus., Q.W., I.L.A., H.A-C., V.A., K.J.A., P.L.A., A.Au., A.B., H.Bec., S.Be., J.Ben., M.Berm., C.Bl., S.E.B., B.Bon., A-L.B-BD., H.Bra., H.Bre., A.B-W., T.B., B.Bur., S.S.B., F.C., J.E.C., J.C-C., S.J.C., G.C-T., C.L.C., NBCS, JM.C., A.Cox., S.S.C., K.Cz., M.B.D., P.D., T.D., M.Dw., D.M.E., DG.E., P.A.F., J.Fi., G.Fl., M.G-D., S.M.G, J.A.G-S., M.M.G., G.G.G., M.S.G., A.G-N., G.I.G, M.Grip., P.Gu., C.A.H., P.Hall., U.H., E.F.H., B.A.M.H-G., B.Ho., A.Hol., M.J.H., R.N.H., J.L.Ho., A.How., kConFab/AOCS, M.Ja., A.Jak., E.M.J., M.E.J., A.Ju., R.Ka., S.Kaup., R.Ke., E.Kh., C.M.Ki., Y-D.K., S.Kou., V.N.K., U.K., K.K-S., A.W.K., K.Ky., D.La., D.G.L., A.Lin., M.Lin., J.Lis., A.L., W-L.L., R.J.M., A.Man., M.Man., S.Mar., ME.M., C.Mc., A.Me., U.Me., H.Ne., W.G.N., J.No., K.Of., H.O., N.O., T-W.P-S., A.V.P., J.Pet., G.Pi., D.P.K., R.P., K.Pu., K.Py., P.Ra., G.R., A.Ro., T.Rü., E.S., S.S., D.P.S., E.J.S., R.K.S., M.J.S., B.Sch., M.E.S., X-O.S., S.Sm., M.C.S., J.J.S., A.J.S., R.M.T., W.J.T., J.A.T., L.R.T., MB.T., D.T., M.A.T., C.M.V, C.H.M.VD., E.M.VV., P.Wa., C.R.W., C.We., J.We., R.Wi., A.W., X.R.Y., W.Z., F.J.C., J.Si., P.Kr., D.F.E., P.D.P.P., M.K.S., M.G-C. All authors read and approved the final version of the manuscript.

## Acknowledgments

A full description of the acknowledgments is provided in the **Supplemental Funding and Acknowledgement**.

