## Supplementary_methods for "Common variants in breast cancer risk loci predispose to distinct tumor subtypes"

**Supplemental Methods**

**Study Population and Genotyping: Supplementary Table 1** provides a description of the studies that were included in these analyses. We included invasive cases and controls from 81 BCAC studies from over 20 countries with genotyping data derived from two Illumina genome-wide custom arrays, the iCOGS and Oncoarray (BCAC data freeze v10 for phenotype data and v10 for genotype/imputation data). Details on the study population, genotype calling, quality control and imputation for the OncoArray and iCOGS are described elsewhere(1-4). Most studies were population-based case–control studies, or case–control studies nested within population-based cohorts, but a subset of studies oversampled cases with a family history of the disease. Information on clinicopathogic characteristics were collected by the individual studies under IRBs and combined in a central database after quality control checks. We only included cases of the first invasive breast cancer, thus we excluded invasive second or contralateral breast cancers and cases diagnosed with carcinoma *in situ* or with missing information on invasiveness. We also excluded all studies from a study specific country if a study country did not contribute data on controls, or if tumor marker data (ER, PR and HER2) was missing on two or more of the tumor marker subtypes (see footnote of **Supplementary Table 1** for further explanation of excluded studies).

We evaluated for evidence of heterogeneity among breast cancer variants that were identified or replicated as being associated with risk of breast cancer in the largest, to date, breast cancer GWAS from the Breast Cancer Association Consortium – 153 variants associated with risk of overall breast cancer risk(1) and 20 variants associated with risk of ER-negative breast cancer(2). We investigated variants that were reported to be associated with risk of breast cancer at p-value threshold p<5.0x10^-8^ (**Supplementary Table 2**)(1,2). Variants were not selected based on prior evidence of having heterogenous associations with breast cancer tumor markers. The investigated variants were excluded in prior analyses that aimed to identify novel breast cancer risk variants that display evidence of tumor heterogeneity(5). Genotyped or imputed SNPs marking each of the 173 susceptibility loci were determined using the iCOGS and the OncoArray genotyping arrays and imputation to the 1000 Genomes Project (Phase 3) reference panel. All imputed SNPs had r^2^>0.5 and the average r^2^ among the selected 173 SNPs in iCOGS and OncoArray was 0.95 and 0.99, respectively. A subset of participants was genotyped on both iCOGS and Oncoarray arrays and for these subjects we only analyzed the Oncoarray data.

**Statistical Analysis:** The two-stage polytomous logistic regression model allows us to efficiently characterize heterogeneity while accounting for tumor marker correlations and large amounts of missing tumor data.(6) The first stage of the model uses a polytomous logistic regression to model case-control odds ratios (ORs) between the susceptibility SNPs and all possible subtypes that could be defined by the combination of tumor markers. For example, in a model fit to evaluate heterogeneity according to ER, PR and HER2 positive/negative status, and grade of differentiation (low, intermediate and high grade), the first stage incorporates case-control ORs for 24 subtypes. The second stage restructures the first-stage subtype-specific case-control ORs parameters into second-stage parameters through a decomposition procedure resulting in a second-stage baseline parameter that represents a case-control OR of a baseline cancer subtype, and case-case ORs parameters for each individual tumor characteristic. The second-stage case-case parameters can be used to perform heterogeneity tests with respect to each specific tumor marker while adjusting for the other tumor markers in the model. The two-stage model efficiently handles missing data by implementing an Expectation-Maximization algorithm (7) that essentially performs iterative “imputation” of the missing tumor characteristics conditional on available tumor characteristics and baseline covariates based on an underlying two-stage polytomous model.

Our criteria for identifying SNPs that showed evidence of heterogeneity consisted of first using the mixed-effects two-stage polytomous model (model 1) to perform the first-stage global heterogeneity test to identify SNPs with evidence of having risk estimates that vary by at least one of the underlying tumor characteristics. In this model, we keep the second stage main effect of ER as a fixed effect since there is strong *a priori* evidence that ER is a common source of heterogeneity(8). As there is less evidence suggesting that PR, HER2, and grade are common sources of heterogeneity, we assume the case-case parameter of PR, HER2 and grade as random effects. We accounted for multiple testing in the global heterogeneity test using a False Discovery Rate (FDR) < 0.05 under the Benjamini-Hochberg procedure(9). We evaluated for potential inflation in the global heterogeneity test due to study/batch effects by permuting genotypes of controls and cases in each study, no evidence of inflation was identified. Second, among SNPs with evidence of global heterogeneity, we used the fixed-effects second-stage specific tumor marker heterogeneity test (model 2) to identify which tumor marker(s) contributed to observed heterogeneity. We considered a specific tumor marker heterogeneity test p-value < 0.05 as evidence of a tumor marker contributing to heterogeneity. We fit a separate fixed-effect two-stage models (model 3) to estimate case-control ORs and 95% confidence intervals (CI) for five surrogate intrinsic-like subtypes defined by combinations of ER, PR, HER2 and grade: (1) luminal A-like (ER+ and/or PR+, HER2-, grade 1 & 2); (2) luminal B-like/HER2-negative (ER+ and/or PR+, HER2-, grade 3); (3) luminal B-like/HER2-positive (ER+ and/or PR+, HER2+); (4) HER2-positive/non-luminal (ER- and PR-, HER2+), and (5) TN (ER-, PR-, HER2-). For the 12 variants associated exclusively with grade in case-case comparisons (model 2), we fit a separate fixed-effect two-stage models (model 4) to estimate case-control ORs and 95% confidence intervals (CI) for tumor grade (grade 1, grade 2, and grade 3).

To help describe sources of heterogeneity, we performed cluster analyses based on Euclidean distance calculated from z-statistics that were estimated by the individual tumor marker heterogeneity tests and by the case-control tests between variants and the intrinsic-like subtypes. Clustering was performed in R using the function Heatmap as implemented by the package “Complex Heatmap” version 2.62(10).

The implementation of this two-stage polytomous regression method is available in a R package called TOP (<https://github.com/andrewhaoyu/TOP>) with a detailed tutorial available at <https://github.com/andrewhaoyu/TOP/blob/master/inst/TOP.pdf>.
