## Supplementary_figures for "Common variants in breast cancer risk loci predispose to distinct tumor subtypes"

**Supplementary Figure 3**. Results from fixed-effects two-stage polytomous models for risk associations^a^ with intrinsic-like subtypes (model 3) for 85 variants with evidence for global heterogeneity^b^. Variants presented in order as shown in **Supplementary** **Figure 2:** **A)** variants with the strongest evidence of associations with luminal-like subtypes, **B)** variants with strong evidence for associations with risk for all subtypes, **C)** variants with stronger evidence for TN or non-luminal subtype association, and **D)** variants predominately associated with luminal-like subtypes.

Odds ratio and 95% CI

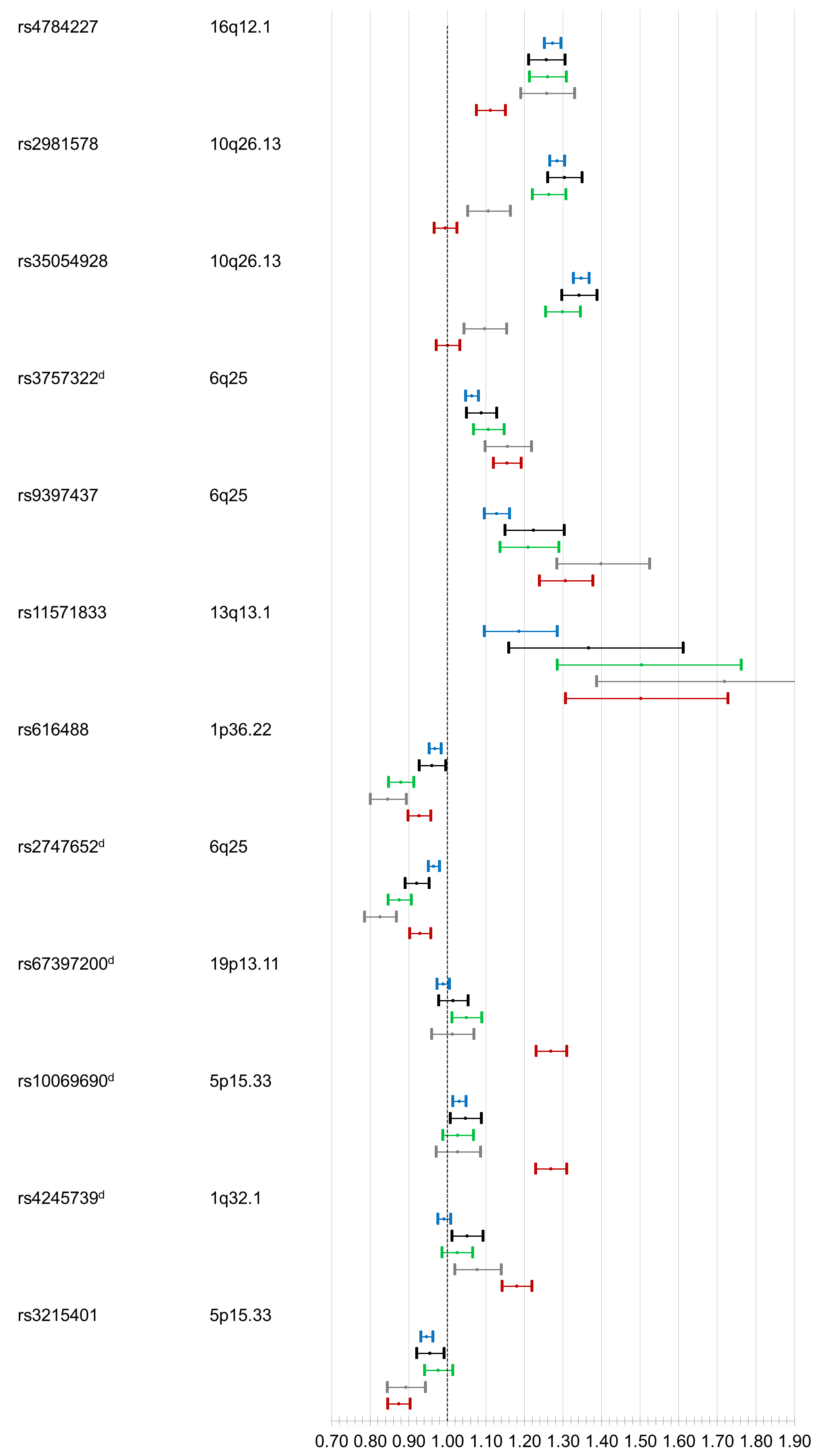

SNP

Locus

Intrinsic-like subtypes^c^

**A**

**B**

**C**

SNP

Locus

Intrinsic-like subtypes^c^

Odds ratio and 95% CI

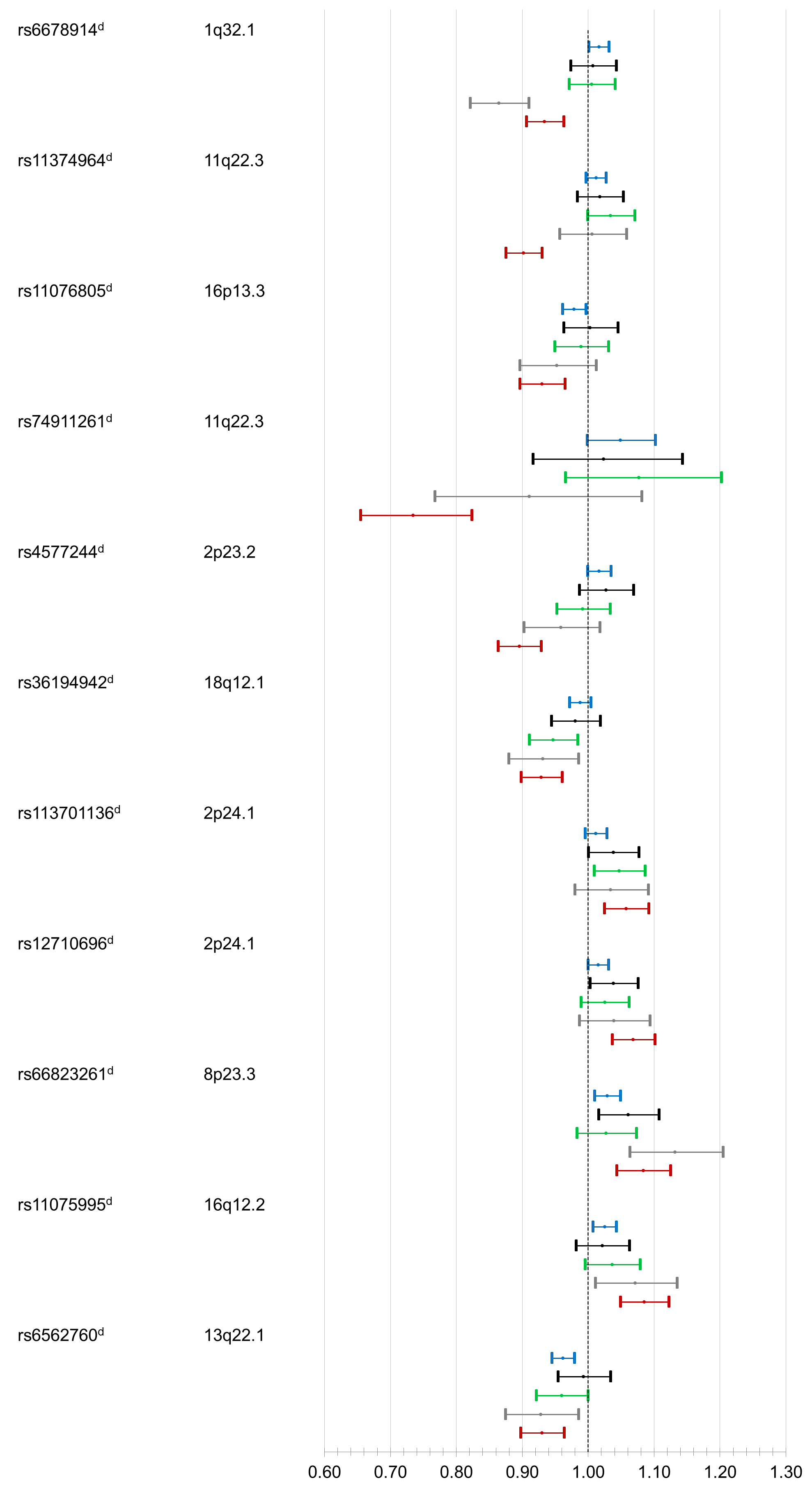

**C cont**

a Per-minor allele odds ratio (95% confidence limits).

^b^ Model 1, Mixed-effect two-stage polytomous model (ER as fixed-effect, and PR, HER2 and grade as random-effects) testing for global heterogeneity between each individual susceptibility variant and any of the tumor features

^c^ Luminal A-like (ER+ and/or PR+, HER2-, grade 1 & 2); Luminal B-like/HER2-negative (ER+ and/or PR+, HER2-, grade 3); luminal B-like/HER2-positive (ER+ and/or PR+, HER2+); HER2-positive/non-luminal (ER- and PR-, HER2+), and triple-negative (ER-, PR-, HER2-)

Luminal A-like	Luminal B-like/HER2-negative	luminal B-like/HER2-positive	HER2-positive/non-luminal	Triple-Negative

**Supplementary Figure 3 cont**. Results from fixed-effects two-stage polytomous models for risk associations^a^ with intrinsic-like subtypes (model 3) for 85 variants with evidence for global heterogeneity^b^. Variants presented in order as shown in **Supplementary** **Figure 2: A)** variants with the strongest evidence of associations with luminal-like subtypes, **B)** variants with strong evidence for associations with risk for all subtypes, **C)** variants with stronger evidence for TN or non-luminal subtype association, and **D)** variants predominately associated with luminal-like subtypes.

Odds ratio and 95% CI

Odds ratio and 95% CI

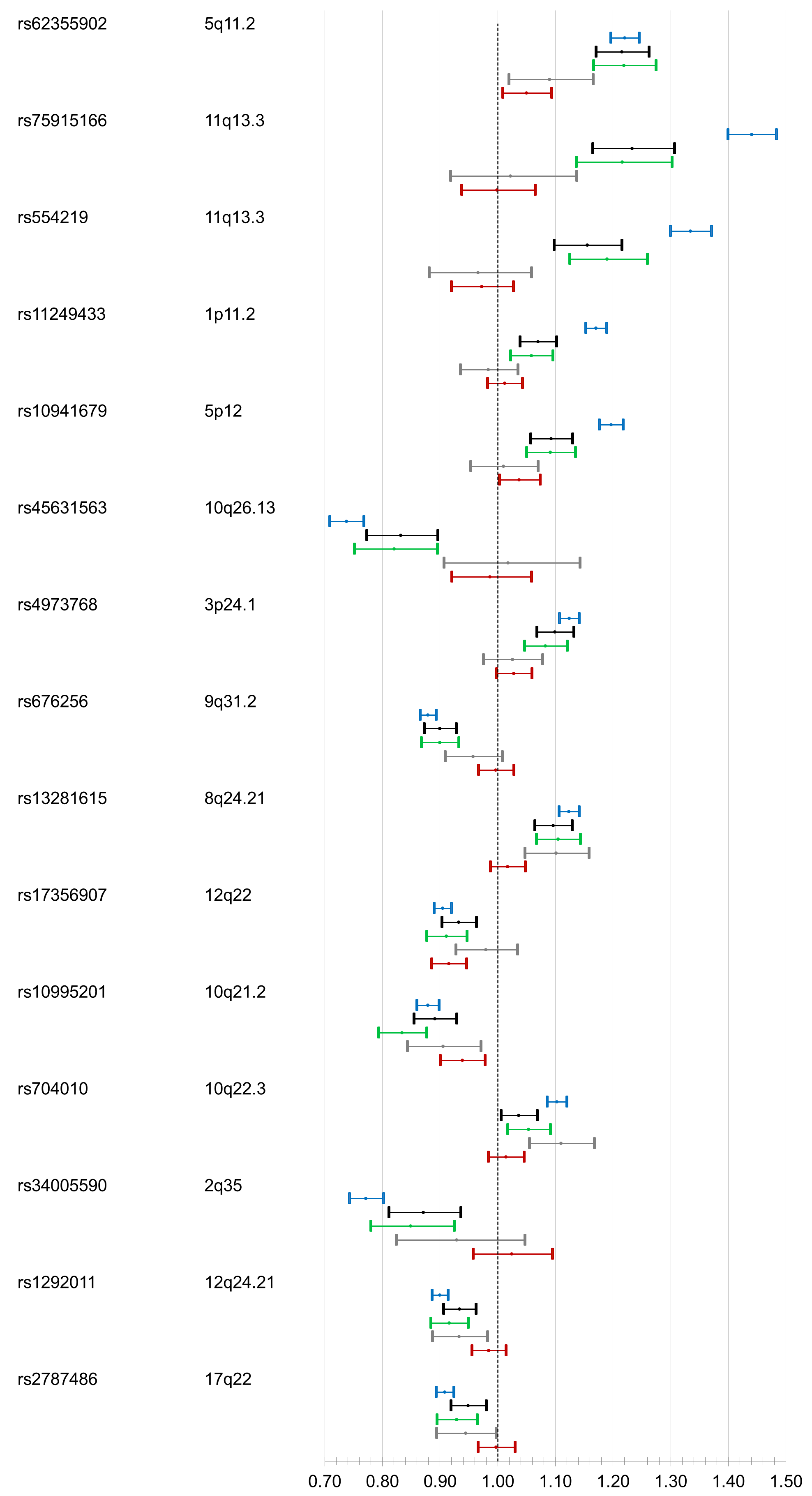

SNP

Locus

Intrinsic-like subtypes^c^

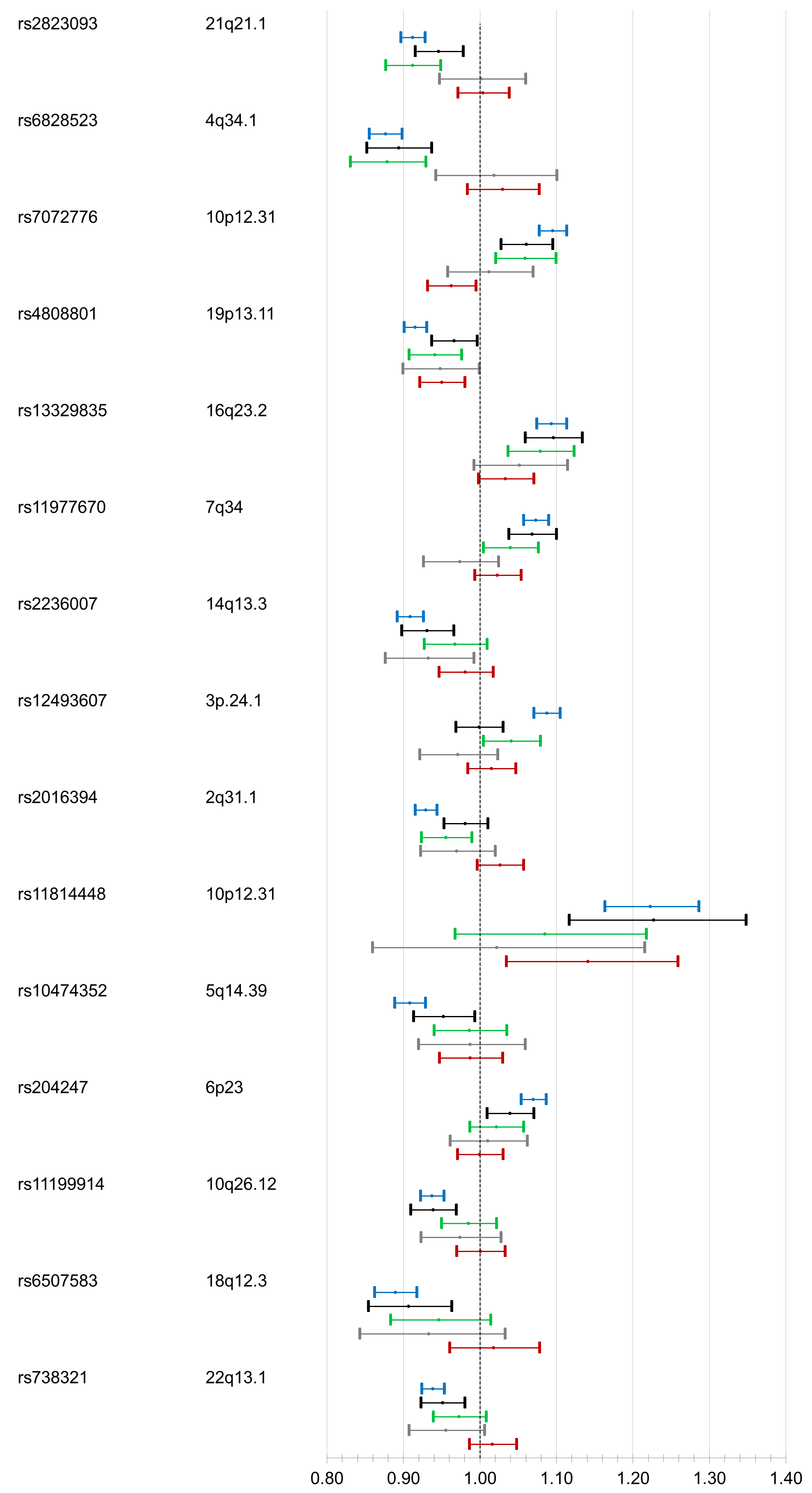

SNP

Locus

Intrinsic-like subtypes^c^

a Per-minor allele odds ratio (95% confidence limits).

^b^ Model 1, Mixed-effect two-stage polytomous model (ER as fixed-effect, and PR, HER2 and grade as random-effects) testing for global heterogeneity between each individual susceptibility variant and any of the tumor features

^c^ Luminal A-like (ER+ and/or PR+, HER2-, grade 1 & 2); Luminal B-like/HER2-negative (ER+ and/or PR+, HER2-, grade 3); luminal B-like/HER2-positive (ER+ and/or PR+, HER2+); HER2-positive/non-luminal (ER- and PR-, HER2+), and triple-negative (ER-, PR-, HER2-)

Luminal A-like	Luminal B-like/HER2-negative	luminal B-like/HER2-positive	HER2-positive/non-luminal	Triple-Negative

**D**

**D cont**

**Supplementary Figure 3 cont**. Results from fixed-effects two-stage polytomous models for risk associations^a^ with intrinsic-like subtypes (model 3) for 85 variants with evidence for global heterogeneity^b^. Variants presented in order as shown in **Supplementary** **Figure 2: A)** variants with the strongest evidence of associations with luminal-like subtypes, **B)** variants with strong evidence for associations with risk for all subtypes, **C)** variants with stronger evidence for TN or non-luminal subtype association, and **D)** variants predominately associated with luminal-like subtypes.

a Per-minor allele odds ratio (95% confidence limits).

^b^ Model 1, Mixed-effect two-stage polytomous model (ER as fixed-effect, and PR, HER2 and grade as random-effects) testing for global heterogeneity between each individual susceptibility variant and any of the tumor features

^c^ Luminal A-like (ER+ and/or PR+, HER2-, grade 1 & 2); Luminal B-like/HER2-negative (ER+ and/or PR+, HER2-, grade 3); luminal B-like/HER2-positive (ER+ and/or PR+, HER2+); HER2-positive/non-luminal (ER- and PR-, HER2+), and triple-negative (ER-, PR-, HER2-)

Luminal A-like	Luminal B-like/HER2-negative	luminal B-like/HER2-positive	HER2-positive/non-luminal	Triple-Negative

Odds ratio and 95% CI

Odds ratio and 95% CI

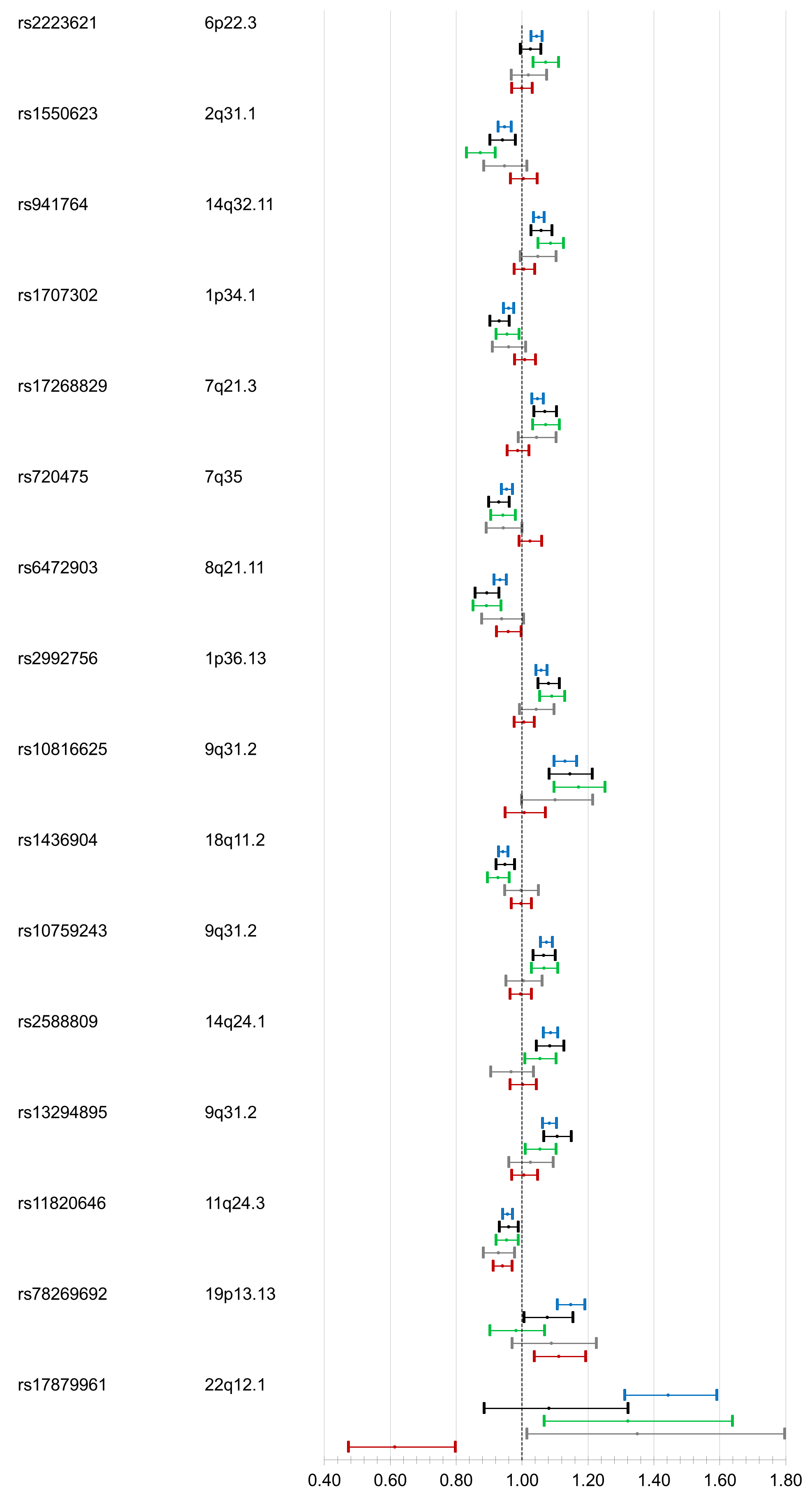

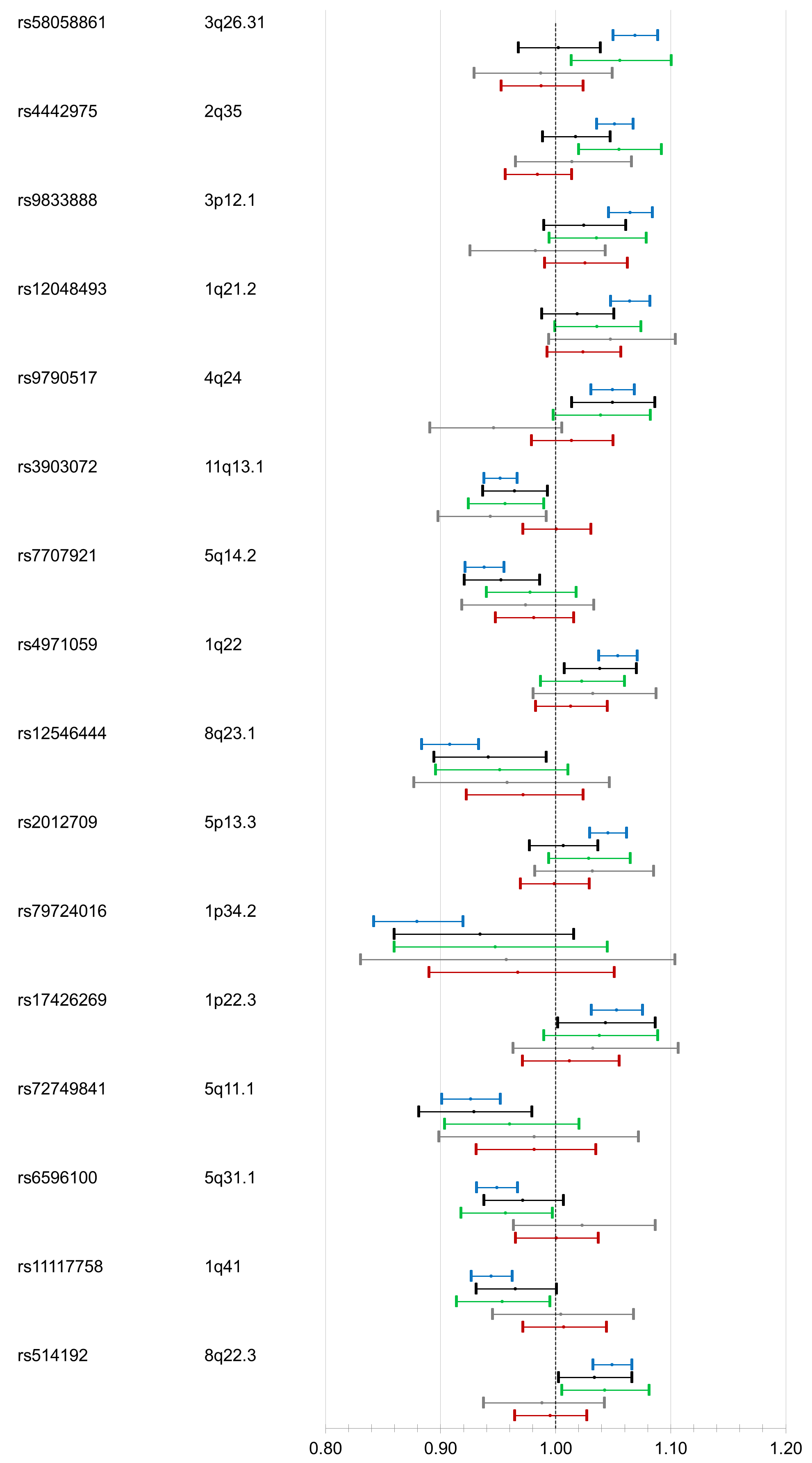

SNP

Locus

Intrinsic-like subtypes

SNP

Locus

Intrinsic-like subtypes

**D cont**

**D**

**Supplementary Figure 4.** Results from fixed-effects two-stage polytomous models for risk associations^a^ with tumor grade (model 4) for the 12 variants associated at P<0.05 only with grade in case-case comparisons (from model 2)^b^

Grade 1	Grade 2	Grade 3

a Per-minor allele odds ratio (95% confidence limits).

^b^ Model 2, Fixed effect two-stage polytomous model for marker-specific tumor heterogeneity tests (i.e. case-case comparisons) between each individual variant and each of the tumor features (ER, PR, HER2, and grade), mutually adjusted for each other

Odds ratio and 95% CI

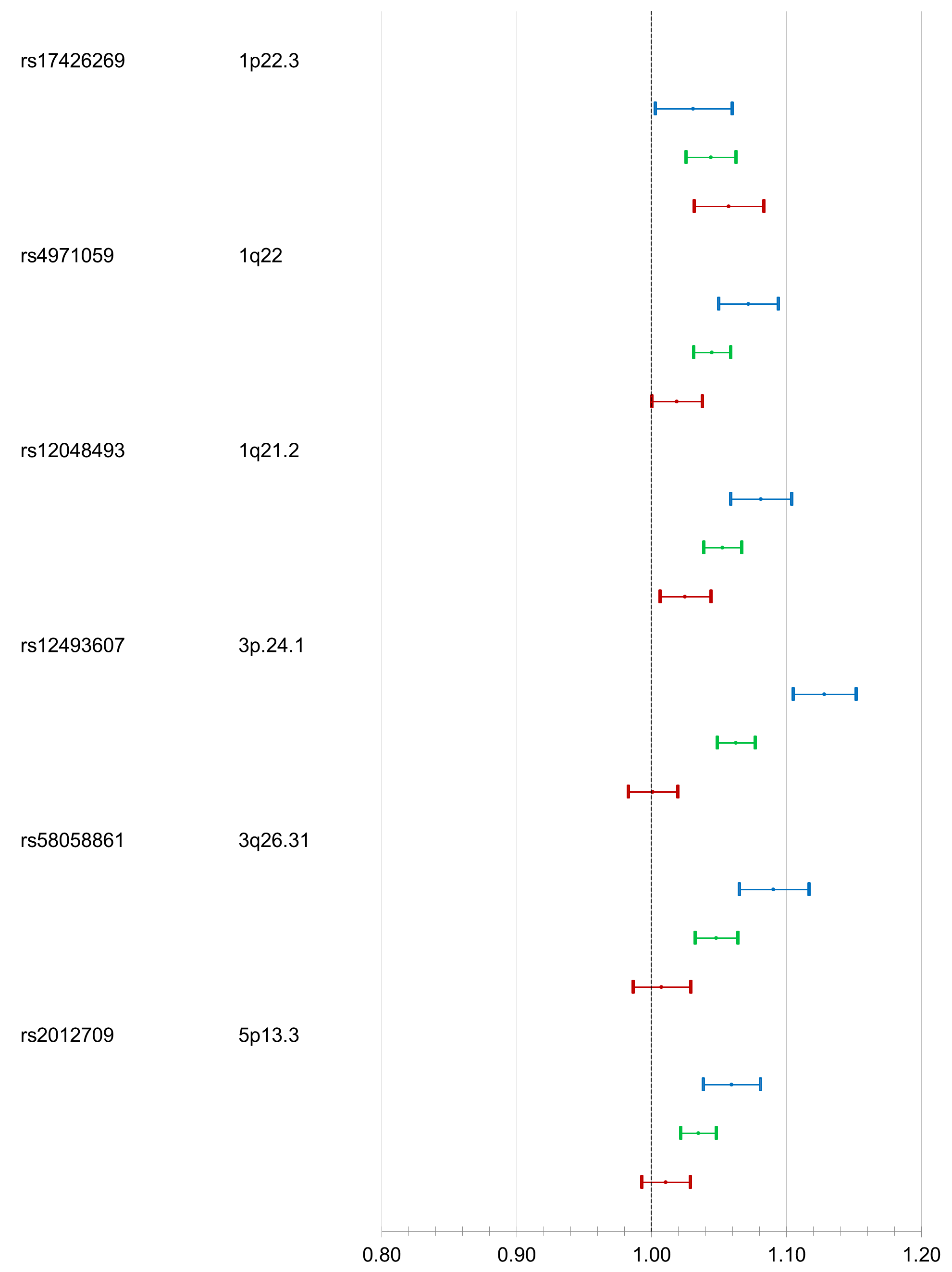

SNP

Locus

Breast cancer risk by subtypes

Odds ratio and 95% CI

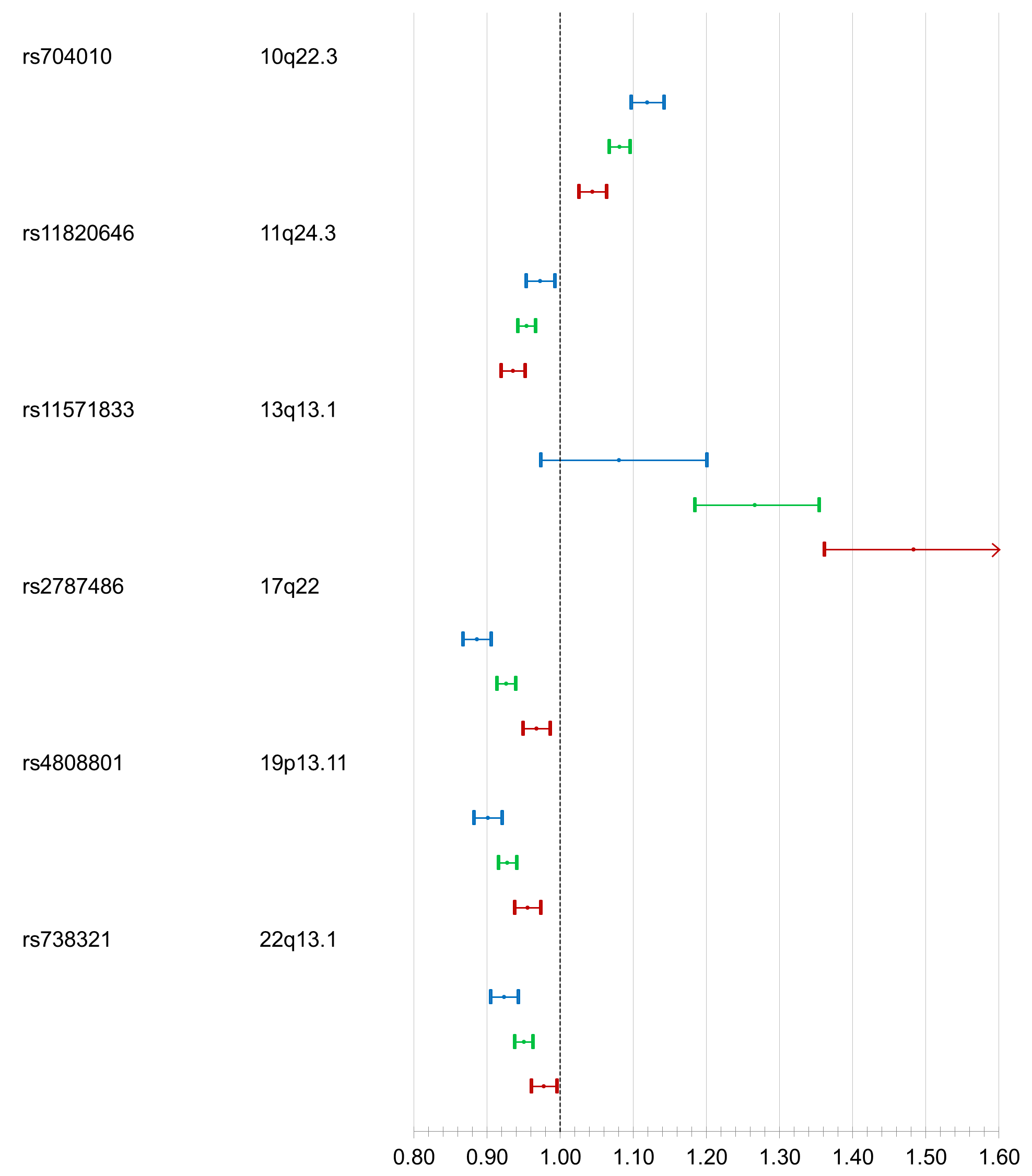

SNP

Locus

Breast cancer risk by subtypes
