## Supplementary material for "Common variants in breast cancer risk loci predispose to distinct tumor subtypes": Fundings and acknowledgement

**Supplemental Funding and Acknowledgments**

**Funding**

This project has been funded in whole or in part with Federal funds from the National Cancer Institute, National Institutes of Health, under NCI Contract No. 75N910D00024. BCAC is funded by Cancer Research UK [C1287/A16563, C1287/A10118], the European Union's Horizon 2020 Research and Innovation Programme (grant numbers 634935 and 633784 for BRIDGES and B-CAST respectively), and by the European Community´s Seventh Framework Programme under grant agreement number 223175 (grant number HEALTH-F2-2009-223175) (COGS). The EU Horizon 2020 Research and Innovation Programme funding source had no role in study design, data collection, data analysis, data interpretation or writing of the report.

Genotyping of the OncoArray was funded by the NIH Grant U19 CA148065, and Cancer UK Grant C1287/A16563 and the PERSPECTIVE project supported by the Government of Canada through Genome Canada and the Canadian Institutes of Health Research (grant GPH-129344) and, the Ministère de l’Économie, Science et Innovation du Québec through Genome Québec and the PSRSIIRI-701 grant, and the Quebec Breast Cancer Foundation. Funding for the iCOGS infrastructure came from: the European Community's Seventh Framework Programme under grant agreement n° 223175 (HEALTH-F2-2009-223175) (COGS), Cancer Research UK (C1287/A10118, C1287/A10710, C12292/A11174, C1281/A12014, C5047/A8384, C5047/A15007, C5047/A10692, C8197/A16565), the National Institutes of Health (CA128978) and Post-Cancer GWAS initiative (1U19 CA148537, 1U19 CA148065 and 1U19 CA148112 - the GAME-ON initiative), the Department of Defence (W81XWH-10-1-0341), the Canadian Institutes of Health Research (CIHR) for the CIHR Team in Familial Risks of Breast Cancer, and Komen Foundation for the Cure, the Breast Cancer Research Foundation, and the Ovarian Cancer Research Fund. The DRIVE Consortium was funded by U19 CA148065.

The Australian Breast Cancer Family Study (ABCFS) was supported by grant UM1 CA164920 from the National Cancer Institute (USA). The content of this manuscript does not necessarily reflect the views or policies of the National Cancer Institute or any of the collaborating centers in the Breast Cancer Family Registry (BCFR), nor does mention of trade names, commercial products, or organizations imply endorsement by the USA Government or the BCFR. The ABCFS was also supported by the National Health and Medical Research Council of Australia, the New South Wales Cancer Council, the Victorian Health Promotion Foundation (Australia) and the Victorian Breast Cancer Research Consortium. J.L.H. is a National Health and Medical Research Council (NHMRC) Senior Principal Research Fellow. M.C.S. is a NHMRC Senior Research Fellow. The ABCS study was supported by the Dutch Cancer Society [grants NKI 2007-3839; 2009 4363]. The Australian Breast Cancer Tissue Bank (ABCTB) was supported by the National Health and Medical Research Council of Australia, The Cancer Institute NSW and the National Breast Cancer Foundation. The AHS study is supported by the intramural research program of the National Institutes of Health, the National Cancer Institute (grant number Z01-CP010119), and the National Institute of Environmental Health Sciences (grant number Z01-ES049030). The work of the BBCC was partly funded by ELAN-Fond of the University Hospital of Erlangen. The BBCS is funded by Cancer Research UK and Breast Cancer Now and acknowledges NHS funding to the NIHR Biomedical Research Centre, and the National Cancer Research Network (NCRN). The BCEES was funded by the National Health and Medical Research Council, Australia and the Cancer Council Western Australia and acknowledges funding from the National Breast Cancer Foundation (JS). For the BCFR-NY, BCFR-PA, BCFR-UT this work was supported by grant UM1 CA164920 from the National Cancer Institute. The content of this manuscript does not necessarily reflect the views or policies of the National Cancer Institute or any of the collaborating centers in the Breast Cancer Family Registry (BCFR), nor does mention of trade names, commercial products, or organizations imply endorsement by the US Government or the BCFR. For BIGGS, ES is supported by NIHR Comprehensive Biomedical Research Centre, Guy's & St. Thomas' NHS Foundation Trust in partnership with King's College London, United Kingdom. IT is supported by the Oxford Biomedical Research Centre. The BREast Oncology GAlician Network (BREOGAN) is funded by Acción Estratégica de Salud del Instituto de Salud Carlos III FIS PI12/02125/Cofinanciado FEDER; Acción Estratégica de Salud del Instituto de Salud Carlos III FIS Intrasalud (PI13/01136); Programa Grupos Emergentes, Cancer Genetics Unit, Instituto de Investigacion Biomedica Galicia Sur. Xerencia de Xestion Integrada de Vigo-SERGAS, Instituto de Salud Carlos III, Spain; Grant 10CSA012E, Consellería de Industria Programa Sectorial de Investigación Aplicada, PEME I + D e I + D Suma del Plan Gallego de Investigación, Desarrollo e Innovación Tecnológica de la Consellería de Industria de la Xunta de Galicia, Spain; Grant EC11-192. Fomento de la Investigación Clínica Independiente, Ministerio de Sanidad, Servicios Sociales e Igualdad, Spain; and Grant FEDER-Innterconecta. Ministerio de Economia y Competitividad, Xunta de Galicia, Spain. The BSUCH study was supported by the Dietmar-Hopp Foundation, the Helmholtz Society and the German Cancer Research Center (DKFZ). CBCS is funded by the Canadian Cancer Society (grant # 313404) and the Canadian Institutes of Health Research. CCGP is supported by funding from the University of Crete. The CECILE study was supported by Fondation de France, Institut National du Cancer (INCa), Ligue Nationale contre le Cancer, Agence Nationale de Sécurité Sanitaire, de l'Alimentation, de l'Environnement et du Travail (ANSES), Agence Nationale de la Recherche (ANR). The CGPS was supported by the Chief Physician Johan Boserup and Lise Boserup Fund, the Danish Medical Research Council, and Herlev and Gentofte Hospital. The CNIO-BCS was supported by the Instituto de Salud Carlos III, the Red Temática de Investigación Cooperativa en Cáncer and grants from the Asociación Española Contra el Cáncer and the Fondo de Investigación Sanitario (PI11/00923 and PI12/00070). Diana Torres was in part supported by a postdoctoral fellowship from the Alexander von Humboldt Foundation. The American Cancer Society funds the creation, maintenance, and updating of the CPS-II cohort. The CTS was initially supported by the California Breast Cancer Act of 1993 and the California Breast Cancer Research Fund (contract 97-10500) and is currently funded through the National Institutes of Health (R01 CA77398, UM1 CA164917, and U01 CA199277). Collection of cancer incidence data was supported by the California Department of Public Health as part of the statewide cancer reporting program mandated by California Health and Safety Code Section 103885. HAC receives support from the Lon V Smith Foundation (LVS39420). The University of Westminster curates the DietCompLyf database funded by Against Breast Cancer Registered Charity No. 1121258 and the NCRN. The coordination of EPIC is financially supported by the European Commission (DG-SANCO) and the International Agency for Research on Cancer. The national cohorts are supported by: Ligue Contre le Cancer, Institut Gustave Roussy, Mutuelle Générale de l’Education Nationale, Institut National de la Santé et de la Recherche Médicale (INSERM) (France); German Cancer Aid, German Cancer Research Center (DKFZ), Federal Ministry of Education and Research (BMBF) (Germany); the Hellenic Health Foundation, the Stavros Niarchos Foundation (Greece); Associazione Italiana per la Ricerca sul Cancro-AIRC-Italy and National Research Council (Italy); Dutch Ministry of Public Health, Welfare and Sports (VWS), Netherlands Cancer Registry (NKR), LK Research Funds, Dutch Prevention Funds, Dutch ZON (Zorg Onderzoek Nederland), World Cancer Research Fund (WCRF), Statistics Netherlands (The Netherlands); Health Research Fund (FIS), PI13/00061 to Granada, PI13/01162 to EPIC-Murcia, Regional Governments of Andalucía, Asturias, Basque Country, Murcia and Navarra, ISCIII RETIC (RD06/0020) (Spain); Cancer Research UK (14136 to EPIC-Norfolk; C570/A16491 and C8221/A19170 to EPIC-Oxford), Medical Research Council (1000143 to EPIC-Norfolk, MR/M012190/1 to EPIC-Oxford) (United Kingdom). The ESTHER study was supported by a grant from the Baden Württemberg Ministry of Science, Research and Arts. Additional cases were recruited in the context of the VERDI study, which was supported by a grant from the German Cancer Aid (Deutsche Krebshilfe). FHRISK is funded from NIHR grant PGfAR 0707-10031. The GC-HBOC (German Consortium of Hereditary Breast and Ovarian Cancer) is supported by the German Cancer Aid (grant no 110837, coordinator: Rita K. Schmutzler, Cologne). This work was also funded by the European Regional Development Fund and Free State of Saxony, Germany (LIFE - Leipzig Research Centre for Civilization Diseases, project numbers 713-241202, 713-241202, 14505/2470, 14575/2470). The GENICA was funded by the Federal Ministry of Education and Research (BMBF) Germany grants 01KW9975/5, 01KW9976/8, 01KW9977/0 and 01KW0114, the Robert Bosch Foundation, Stuttgart, Deutsches Krebsforschungszentrum (DKFZ), Heidelberg, the Institute for Prevention and Occupational Medicine of the German Social Accident Insurance, Institute of the Ruhr University Bochum (IPA), Bochum, as well as the Department of Internal Medicine, Evangelische Kliniken Bonn gGmbH, Johanniter Krankenhaus, Bonn, Germany. The GEPARSIXTO study was conducted by the German Breast Group GmbH. The GESBC was supported by the Deutsche Krebshilfe e. V. [70492] and the German Cancer Research Center (DKFZ). The HABCS study was supported by the Claudia von Schilling Foundation for Breast Cancer Research, by the Lower Saxonian Cancer Society, and by the Rudolf Bartling Foundation. The HEBCS was financially supported by the Helsinki University Hospital Research Fund, the Finnish Cancer Society, and the Sigrid Juselius Foundation. The HUBCS was supported by a grant from the German Federal Ministry of Research and Education (RUS08/017), and by the Russian Foundation for Basic Research and the Federal Agency for Scientific Organizations for support the Bioresource collections and RFBR grants 14-04-97088, 17-29-06014 and 17-44-020498. Financial support for KARBAC was provided through the regional agreement on medical training and clinical research (ALF) between Stockholm County Council and Karolinska Institutet, the Swedish Cancer Society, The Gustav V Jubilee foundation and Bert von Kantzows foundation. The KARMA study was supported by Märit and Hans Rausings Initiative Against Breast Cancer. The KBCP was financially supported by the special Government Funding (EVO) of Kuopio University Hospital grants, Cancer Fund of North Savo, the Finnish Cancer Organizations, and by the strategic funding of the University of Eastern Finland. kConFab is supported by a grant from the National Breast Cancer Foundation, and previously by the National Health and Medical Research Council (NHMRC), the Queensland Cancer Fund, the Cancer Councils of New South Wales, Victoria, Tasmania and South Australia, and the Cancer Foundation of Western Australia. Financial support for the AOCS was provided by the United States Army Medical Research and Materiel Command [DAMD17-01-1-0729], Cancer Council Victoria, Queensland Cancer Fund, Cancer Council New South Wales, Cancer Council South Australia, The Cancer Foundation of Western Australia, Cancer Council Tasmania and the National Health and Medical Research Council of Australia (NHMRC; 400413, 400281, 199600). G.C.T. and P.W. are supported by the NHMRC. RB was a Cancer Institute NSW Clinical Research Fellow. The CSP is also part of the National Cancer Institute's Division of Cancer Prevention and Control Surveillance, Epidemiology, and End Results Program, under contract number N01CN25403. DL is supported by the FWO. The MABCS study is funded by the Research Centre for Genetic Engineering and Biotechnology "Georgi D. Efremov", MASA. The MARIE study was supported by the Deutsche Krebshilfe e.V. [70-2892-BR I, 106332, 108253, 108419, 110826, 110828], the Hamburg Cancer Society, the German Cancer Research Center (DKFZ) and the Federal Ministry of Education and Research (BMBF) Germany [01KH0402]. MBCSG is supported by grants from the Italian Association for Cancer Research (AIRC) and by funds from the Italian citizens who allocated the 5/1000 share of their tax payment in support of the Fondazione IRCCS Istituto Nazionale Tumori, according to Italian laws (INT-Institutional strategic projects “5x1000”). The MCBCS was supported by the NIH grants CA192393, CA116167, CA176785 an NIH Specialized Program of Research Excellence (SPORE) in Breast Cancer [CA116201], and the Breast Cancer Research Foundation and a generous gift from the David F. and Margaret T. Grohne Family Foundation. The Melbourne Collaborative Cohort Study (MCCS) cohort recruitment was funded by VicHealth and Cancer Council Victoria. The MCCS was further augmented by Australian National Health and Medical Research Council grants 209057, 396414 and 1074383 and by infrastructure provided by Cancer Council Victoria. Cases and their vital status were ascertained through the Victorian Cancer Registry and the Australian Institute of Health and Welfare, including the National Death Index and the Australian Cancer Database. The MEC was support by NIH grants CA63464, CA54281, CA098758, CA132839 and CA164973. The MISS study is supported by funding from ERC-2011-294576 Advanced grant, Swedish Cancer Society, Swedish Research Council, Local hospital funds, Berta Kamprad Foundation, Gunnar Nilsson. The MMHS study was supported by NIH grants CA97396, CA128931, CA116201, CA140286 and CA177150. MSKCC is supported by grants from the Breast Cancer Research Foundation and Robert and Kate Niehaus Clinical Cancer Genetics Initiative. The work of MTLGEBCS was supported by the Quebec Breast Cancer Foundation, the Canadian Institutes of Health Research for the “CIHR Team in Familial Risks of Breast Cancer” program – grant # CRN-87521 and the Ministry of Economic Development, Innovation and Export Trade – grant # PSR-SIIRI-701. The NBCS has received funding from the K.G. Jebsen Centre for Breast Cancer Research; the Research Council of Norway grant 193387/V50 (to A-L Børresen-Dale and V.N. Kristensen) and grant 193387/H10 (to A-L Børresen-Dale and V.N. Kristensen), South Eastern Norway Health Authority (grant 39346 to A-L Børresen-Dale) and the Norwegian Cancer Society (to A-L Børresen-Dale and V.N. Kristensen). The NBHS was supported by NIH grant R01CA100374. Biological sample preparation was conducted the Survey and Biospecimen Shared Resource, which is supported by P30 CA68485. The Northern California Breast Cancer Family Registry (NC-BCFR) and Ontario Familial Breast Cancer Registry (OFBCR) were supported by grant UM1 CA164920 from the National Cancer Institute (USA). The content of this manuscript does not necessarily reflect the views or policies of the National Cancer Institute or any of the collaborating centers in the Breast Cancer Family Registry (BCFR), nor does mention of trade names, commercial products, or organizations imply endorsement by the USA Government or the BCFR. The Carolina Breast Cancer Study was funded by Komen Foundation, the National Cancer Institute (P50 CA058223, U54 CA156733, U01 CA179715), and the North Carolina University Cancer Research Fund. The NHS was supported by NIH grants P01 CA87969, UM1 CA186107, and U19 CA148065. The NHS2 was supported by NIH grants UM1 CA176726 and U19 CA148065. The OBCS was supported by research grants from the Finnish Cancer Foundation, the Academy of Finland (grant number 250083, 122715 and Center of Excellence grant number 251314), the Finnish Cancer Foundation, the Sigrid Juselius Foundation, the University of Oulu, the University of Oulu Support Foundation and the special Governmental EVO funds for Oulu University Hospital-based research activities. The ORIGO study was supported by the Dutch Cancer Society (RUL 1997-1505) and the Biobanking and Biomolecular Resources Research Infrastructure (BBMRI-NL CP16). The PBCS was funded by Intramural Research Funds of the National Cancer Institute, Department of Health and Human Services, USA. Genotyping for PLCO was supported by the Intramural Research Program of the National Institutes of Health, NCI, Division of Cancer Epidemiology and Genetics. The PLCO is supported by the Intramural Research Program of the Division of Cancer Epidemiology and Genetics and supported by contracts from the Division of Cancer Prevention, National Cancer Institute, National Institutes of Health. The POSH study is funded by Cancer Research UK (grants C1275/A11699, C1275/C22524, C1275/A19187, C1275/A15956 and Breast Cancer Campaign 2010PR62, 2013PR044. PROCAS is funded from NIHR grant PGfAR 0707-10031. The RBCS was funded by the Dutch Cancer Society (DDHK 2004-3124, DDHK 2009-4318). The SASBAC study was supported by funding from the Agency for Science, Technology and Research of Singapore (A*STAR), the US National Institute of Health (NIH) and the Susan G. Komen Breast Cancer Foundation. The SBCS was supported by Sheffield Experimental Cancer Medicine Centre and Breast Cancer Now Tissue Bank. SEARCH is funded by Cancer Research UK [C490/A10124, C490/A16561] and supported by the UK National Institute for Health Research Biomedical Research Centre at the University of Cambridge. The University of Cambridge has received salary support for PDPP from the NHS in the East of England through the Clinical Academic Reserve. The Sister Study (SISTER) is supported by the Intramural Research Program of the NIH, National Institute of Environmental Health Sciences (Z01-ES044005 and Z01-ES049033). The Two Sister Study (2SISTER) was supported by the Intramural Research Program of the NIH, National Institute of Environmental Health Sciences (Z01-ES044005 and Z01-ES102245), and, also by a grant from Susan G. Komen for the Cure, grant FAS0703856. SKKDKFZS is supported by the DKFZ. The SMC is funded by the Swedish Cancer Foundation. The SZBCS was supported by Grant PBZ_KBN_122/P05/2004. The TBCS was funded by The National Cancer Institute Thailand. The TNBCC was supported by: a Specialized Program of Research Excellence (SPORE) in Breast Cancer (CA116201), a grant from the Breast Cancer Research Foundation, a generous gift from the David F. and Margaret T. Grohne Family Foundation. The UCIBCS component of this research was supported by the NIH [CA58860, CA92044] and the Lon V Smith Foundation [LVS39420]. The UKBGS is funded by Breast Cancer Now and the Institute of Cancer Research (ICR), London. ICR acknowledges NHS funding to the NIHR Biomedical Research Centre. The UKOPS study was funded by The Eve Appeal (The Oak Foundation) with investigators and supported by the National Institute for Health Research University College London Hospitals Biomedical Research Centre and MRC core funding (MR_UU_12023). The US3SS study was supported by Massachusetts (K.M.E., R01CA47305), Wisconsin (P.A.N., R01 CA47147) and New Hampshire (L.T.-E., R01CA69664) centers, and Intramural Research Funds of the National Cancer Institute, Department of Health and Human Services, USA. The USRT Study was funded by Intramural Research Funds of the National Cancer Institute, Department of Health and Human Services, USA. The WHI program is funded by the National Heart, Lung, and Blood Institute, the US National Institutes of Health and the US Department of Health and Human Services (HHSN268201100046C, HHSN268201100001C, HHSN268201100002C, HHSN268201100003C, HHSN268201100004C and HHSN271201100004C). Dr. Nilanjan Chatterjee was supported by NHGRI (1R01 HG010480-01). This work was also funded by NCI U19 CA148065-01.

**Acknowledgements**

We thank all the individuals who took part in these studies and all the researchers, clinicians, technicians and administrative staff who have enabled this work to be carried out. The COGS study would not have been possible without the contributions of the following: Andrew Berchuck (OCAC), Rosalind A. Eeles, Ali Amin Al Olama, Zsofia  Kote-Jarai, Sara Benlloch (PRACTICAL), Antonis Antoniou, Lesley McGuffog, Andrew Lee, Ed Dicks, Craig Luccarini and the staff of the Centre for Genetic Epidemiology Laboratory, Anna Gonzalez-Neira and the staff of the CNIO genotyping unit, Daniel C. Tessier, Francois Bacot, Daniel Vincent, Sylvie LaBoissière and Frederic Robidoux and the staff of the McGill University and Génome Québec Innovation Centre, Sune F. Nielsen, Borge G. Nordestgaard, and the staff of the Copenhagen DNA laboratory, and Julie M. Cunningham, Sharon A. Windebank, Christopher A. Hilker, Jeffrey Meyer and the staff of Mayo Clinic Genotyping Core Facility. ABCFS thank Maggie Angelakos, Judi Maskiell, Gillian Dite. ABCS thanks the Blood bank Sanquin, The Netherlands. ABCTB Investigators: Christine Clarke, Rosemary Balleine, Robert Baxter, Stephen Braye, Jane Carpenter, Jane Dahlstrom, John Forbes, Soon Lee, Debbie Marsh, Adrienne Morey, Nirmala Pathmanathan, Rodney Scott, Allan Spigelman, Nicholas Wilcken, Desmond Yip. Samples are made available to researchers on a non-exclusive basis. BBCS thanks Eileen Williams, Elaine Ryder-Mills, Kara Sargus. BCEES thanks Allyson Thomson, Christobel Saunders, Terry Slevin, BreastScreen Western Australia, Elizabeth Wylie, Rachel Lloyd. The BCINIS study would not have been possible without the contributions of Dr. K. Landsman, Dr. N. Gronich, Dr. A. Flugelman, Dr. W. Saliba, Dr. E. Liani, Dr. I. Cohen, Dr. S. Kalet, Dr. V. Friedman, Dr. O. Barnet of the NICCC in Haifa, and all the contributing family medicine, surgery, pathology and oncology teams in all medical institutes in Northern Israel. BIGGS thanks Niall McInerney, Gabrielle Colleran, Andrew Rowan, Angela Jones. The BREOGAN study would not have been possible without the contributions of the following: Manuela Gago-Dominguez, Jose Esteban Castelao, Angel Carracedo, Victor Muñoz Garzón, Alejandro Novo Domínguez, Maria Elena Martinez, Sara Miranda Ponte, Carmen Redondo Marey, Maite Peña Fernández, Manuel Enguix Castelo, Maria Torres, Manuel Calaza (BREOGAN), José Antúnez, Máximo Fraga and the staff of the Department of Pathology and Biobank of the University Hospital Complex of Santiago-CHUS, Instituto de Investigación Sanitaria de Santiago, IDIS, Xerencia de Xestion Integrada de Santiago-SERGAS; Joaquín González-Carreró and the staff of the Department of Pathology and Biobank of University Hospital Complex of Vigo, Instituto de Investigacion Biomedica Galicia Sur, SERGAS, Vigo, Spain. BSUCH thanks Peter Bugert, Medical Faculty Mannheim. CBCS thanks study participants, co-investigators, collaborators and staff of the Canadian Breast Cancer Study, and project coordinators Agnes Lai and Celine Morissette. CCGP thanks Styliani Apostolaki, Anna Margiolaki, Georgios Nintos, Maria Perraki, Georgia Saloustrou, Georgia Sevastaki, Konstantinos Pompodakis. CGPS thanks staff and participants of the Copenhagen General Population Study. For the excellent technical assistance: Dorthe Uldall Andersen, Maria Birna Arnadottir, Anne Bank, Dorthe Kjeldgård Hansen. The Danish Cancer Biobank is acknowledged for providing infrastructure for the collection of blood samples for the cases. CNIO-BCS thanks Guillermo Pita, Charo Alonso, Nuria Álvarez, Pilar Zamora, Primitiva Menendez, the Human Genotyping-CEGEN Unit (CNIO). Investigators from the CPS-II cohort thank the participants and Study Management Group for their invaluable contributions to this research. They also acknowledge the contribution to this study from central cancer registries supported through the Centers for Disease Control and Prevention National Program of Cancer Registries, as well as cancer registries supported by the National Cancer Institute Surveillance Epidemiology and End Results program. The CTS Steering Committee includes Leslie Bernstein, Susan Neuhausen, James Lacey, Sophia Wang, Huiyan Ma, and Jessica Clague DeHart at the Beckman Research Institute of City of Hope, Dennis Deapen, Rich Pinder, and Eunjung Lee at the University of Southern California, Pam Horn-Ross, Peggy Reynolds, Christina Clarke Dur and David Nelson at the Cancer Prevention Institute of California, Hoda Anton-Culver, Argyrios Ziogas, and Hannah Park at the University of California Irvine, and Fred Schumacher at Case Western University. DIETCOMPLYF thanks the patients, nurses and clinical staff involved in the study. The DietCompLyf study was funded by the charity Against Breast Cancer (Registered Charity Number 1121258) and the NCRN. We thank the participants and the investigators of EPIC (European Prospective Investigation into Cancer and Nutrition). ESTHER thanks Hartwig Ziegler, Sonja Wolf, Volker Hermann, Christa Stegmaier, Katja Butterbach. FHRISK thanks NIHR for funding. GC-HBOC thanks Stefanie Engert, Heide Hellebrand, Sandra Kröber and LIFE - Leipzig Research Centre for Civilization Diseases (Markus Loeffler, Joachim Thiery, Matthias Nüchter, Ronny Baber). The GENICA Network: Dr. Margarete Fischer-Bosch-Institute of Clinical Pharmacology, Stuttgart, and University of Tübingen, Germany [HB, WYL], German Cancer Consortium (DKTK) and German Cancer Research Center (DKFZ) [HB], Deutsche Forschungsgemeinschaft (DFG, German Research Foundation) under Germany's Excellence Strategy - EXC 2180 - 390900677 [HB], Department of Internal Medicine, Evangelische Kliniken Bonn gGmbH, Johanniter Krankenhaus, Bonn, Germany [YDK, Christian Baisch], Institute of Pathology, University of Bonn, Germany [Hans-Peter Fischer], Molecular Genetics of Breast Cancer, Deutsches Krebsforschungszentrum (DKFZ), Heidelberg, Germany [Ute Hamann], Institute for Prevention and Occupational Medicine of the German Social Accident Insurance, Institute of the Ruhr University Bochum (IPA), Bochum, Germany [TB, Beate Pesch, Sylvia Rabstein, Anne Lotz]; and Institute of Occupational Medicine and Maritime Medicine, University Medical Center Hamburg-Eppendorf, Germany [Volker Harth]. HABCS thanks Michael Bremer. HEBCS thanks Sofia Khan, Johanna Kiiski, Kristiina Aittomäki, Irja Erkkilä. HMBCS thanks Peter Hillemanns, Hans Christiansen and Johann H. Karstens. HUBCS thanks Shamil Gantsev. KARMA and SASBAC thank the Swedish Medical Research Counsel. KBCP thanks Eija Myöhänen, Helena Kemiläinen. kConFab/AOCS wish to thank Heather Thorne, Eveline Niedermayr, all the kConFab research nurses and staff, the heads and staff of the Family Cancer Clinics, and the Clinical Follow Up Study (which has received funding from the NHMRC, the National Breast Cancer Foundation, Cancer Australia, and the National Institute of Health (USA)) for their contributions to this resource, and the many families who contribute to kConFab. MABCS thanks Emilija Lazarova (Clinic of Radiotherapy and Oncology), Dzengis Jasar, Mitko Karadjozov (Adzibadem-Sistina” Hospital), Andrej Arsovski and Liljana Stojanovska (Re-Medika” Hospital) for their contributions and commitment to this study. MARIE thanks Petra Seibold, Dieter Flesch-Janys, Judith Heinz, Nadia Obi, Alina Vrieling, Sabine Behrens, Ursula Eilber, Muhabbet Celik, Til Olchers and Stefan Nickels. MBCSG (Milan Breast Cancer Study Group): Paolo Peterlongo, Siranoush Manoukian, Bernard Peissel, Roberto Villa, Cristina Zanzottera, Bernardo Bonanni, Irene Feroce, and the personnel of the Cogentech Cancer Genetic Test Laboratory. The MCCS was made possible by the contribution of many people, including the original investigators, the teams that recruited the participants and continue working on follow-up, and the many thousands of Melbourne residents who continue to participate in the study. We thank the coordinators, the research staff and especially the MMHS participants for their continued collaboration on research studies in breast cancer. MSKCC thanks Marina Corines, Lauren Jacobs. MTLGEBCS would like to thank Martine Tranchant (CHU de Québec Research Center), Marie-France Valois, Annie Turgeon and Lea Heguy (McGill University Health Center, Royal Victoria Hospital; McGill University) for DNA extraction, sample management and skillful technical assistance. J.S. is Chairholder of the Canada Research Chair in Oncogenetics. The following are NBCS Collaborators: Kristine K. Sahlberg (PhD), Lars Ottestad (MD), Rolf Kåresen (Prof. Em.) Dr. Ellen Schlichting (MD), Marit Muri Holmen (MD), Toril Sauer (MD), Vilde Haakensen (MD), Olav Engebråten (MD), Bjørn Naume (MD), Alexander Fosså (MD), Cecile E. Kiserud (MD), Kristin V. Reinertsen (MD), Åslaug Helland (MD), Margit Riis (MD), Jürgen Geisler (MD), and OSBREAC. NBHS thank study participants and research staff for their contributions and commitment to the studies. For NHS and NHS2 the study protocol was approved by the institutional review boards of the Brigham and Women’s Hospital and Harvard T.H. Chan School of Public Health, and those of participating registries as required. We would like to thank the participants and staff of the NHS and NHS2 for their valuable contributions as well as the following state cancer registries for their help: AL, AZ, AR, CA, CO, CT, DE, FL, GA, ID, IL, IN, IA, KY, LA, ME, MD, MA, MI, NE, NH, NJ, NY, NC, ND, OH, OK, OR, PA, RI, SC, TN, TX, VA, WA, WY. The authors assume full responsibility for analyses and interpretation of these data. OBCS thanks Arja Jukkola-Vuorinen, Mervi Grip, Saila Kauppila, Meeri Otsukka, Leena Keskitalo and Kari Mononen for their contributions to this study. OFBCR thanks Teresa Selander, Nayana Weerasooriya. ORIGO thanks E. Krol-Warmerdam, and J. Blom for patient accrual, administering questionnaires, and managing clinical information. The LUMC survival data were retrieved from the Leiden hospital-based cancer registry system (ONCDOC) with the help of Dr. J. Molenaar. PBCS thanks Louise Brinton, Mark Sherman, Neonila Szeszenia-Dabrowska, Beata Peplonska, Witold Zatonski, Pei Chao, Michael Stagner. The ethical approval for the POSH study is MREC /00/6/69, UKCRN ID: 1137. We thank staff in the Experimental Cancer Medicine Centre (ECMC) supported Faculty of Medicine Tissue Bank and the Faculty of Medicine DNA Banking resource. PREFACE thanks Sonja Oeser and Silke Landrith. PROCAS thanks NIHR for funding. RBCS thanks Petra Bos, Jannet Blom, Ellen Crepin, Elisabeth Huijskens, Anja Kromwijk-Nieuwlaat, Annette Heemskerk, the Erasmus MC Family Cancer Clinic. SBCS thanks Sue Higham, Helen Cramp, Dan Connley, Ian Brock, Sabapathy Balasubramanian and Malcolm W.R. Reed. We thank the SEARCH and EPIC teams. SKKDKFZS thanks all study participants, clinicians, family doctors, researchers and technicians for their contributions and commitment to this study. We thank the SUCCESS Study teams in Munich, Duessldorf, Erlangen and Ulm. SZBCS thanks Ewa Putresza. UCIBCS thanks Irene Masunaka. UKBGS thanks Breast Cancer Now and the Institute of Cancer Research for support and funding of the Breakthrough Generations Study, and the study participants, study staff, and the doctors, nurses and other health care providers and health information sources who have contributed to the study. We acknowledge NHS funding to the Royal Marsden/ICR NIHR Biomedical Research Centre. We acknowledge funding to the NIHR Biomedical Research Centre Manchester (IS-BRC-1215-20007). The authors thank the WHI investigators and staff for their dedication and the study participants for making the program possible.

Support for title page creation and format was provided by AuthorArranger, a tool developed at the National Cancer Institute

**Members of consortia listed as authors**

**NBCS Collaborators**

Kristine K. Sahlberg (Department of Cancer Genetics, Institute for Cancer Research, Oslo University Hospital-Radiumhospitalet, Oslo, Norway; Department of Research, Vestre Viken Hospital, Drammen, Norway), Lars Ottestad (Department of Cancer Genetics, Institute for Cancer Research, Oslo University Hospital-Radiumhospitalet, Oslo, Norway), Rolf Kåresen (Institute of Clinical Medicine, Faculty of Medicine, University of Oslo, Norway; Section for Breast- and Endocrine Surgery, Department of Cancer, Division of Surgery, Cancer and Transplantation Medicine, Oslo University Hospital-Ullevål, Oslo, Norway), Dr. Ellen Schlichting (Section for Breast- and Endocrine Surgery, Department of Cancer, Division of Surgery, Cancer and Transplantation Medicine, Oslo University Hospital-Ullevål, Oslo, Norway), Marit Muri Holmen (Department of Radiology and Nuclear Medicine, Oslo University Hospital, Oslo, Norway), Toril Sauer (Institute of Clinical Medicine, Faculty of Medicine, University of Oslo, Norway; Department of Pathology at Akershus University hospital, Lørenskog, Norway), Vilde Haakensen (Department of Cancer Genetics, Institute for Cancer Research, Oslo University Hospital-Radiumhospitalet, Oslo, Norway), Olav Engebråten (Institute of Clinical Medicine, Faculty of Medicine, University of Oslo, Norway; Department of Tumor Biology, Institute for Cancer Research, Oslo University Hospital, Oslo, Norway; Department of Oncology, Division of Surgery and Cancer and Transplantation Medicine, Oslo University Hospital-Radiumhospitalet, Oslo, Norway), Bjørn Naume (Institute of Clinical Medicine, Faculty of Medicine, University of Oslo, Norway; Department of Oncology, Division of Surgery and Cancer and Transplantation Medicine, Oslo University Hospital-Radiumhospitalet, Oslo, Norway), Alexander Fosså (Department of Oncology, Division of Surgery and Cancer and Transplantation Medicine, Oslo University Hospital-Radiumhospitalet, Oslo, Norway; National Advisory Unit on Late Effects after Cancer Treatment, Department of Oncology, Oslo University Hospital, Oslo, Norway), Cecile E. Kiserud (Department of Oncology, Division of Surgery and Cancer and Transplantation Medicine, Oslo University Hospital-Radiumhospitalet, Oslo, Norway; National Advisory Unit on Late Effects after Cancer Treatment, Department of Oncology, Oslo University Hospital, Oslo, Norway), Kristin V. Reinertsen (Department of Oncology, Division of Surgery and Cancer and Transplantation Medicine, Oslo University Hospital-Radiumhospitalet, Oslo, Norway; National Advisory Unit on Late Effects after Cancer Treatment, Department of Oncology, Oslo University Hospital, Oslo, Norway), Åslaug Helland (Department of Cancer Genetics, Institute for Cancer Research, Oslo University Hospital-Radiumhospitalet, Oslo, Norway; National Advisory Unit on Late Effects after Cancer Treatment, Department of Oncology, Oslo University Hospital, Oslo, Norway), Margit Riis (Section for Breast- and Endocrine Surgery, Department of Cancer, Division of Surgery, Cancer and Transplantation Medicine, Oslo University Hospital-Ullevål, Oslo, Norway), Jürgen Geisler (Institute of Clinical Medicine, Faculty of Medicine, University of Oslo, Norway; Department of Oncology, Akershus University Hospital, Lørenskog, Norway), and OSBREAC (Breast Cancer Research Consortium (Chair: Kristine K. Sahlberg), Oslo University Hospital, Oslo, Norway).

**kConFab/AOCS Investigators**

Stephen Fox, Ian Campbell (Peter MacCallum Cancer Centre, Melbourne, Australia); Georgia Chenevix-Trench, Amanda Spurdle, Penny Webb (QIMR Berghofer Medical Research Institute, Brisbane, Australia); Anna de Fazio (Westmead Millenium Institute, Sydney, Australia); Margaret Tassell (BCNA delegate, Community Representative); Judy Kirk (Westmead Hospital, Sydney, Australia); Geoff Lindeman (Walter and Eliza

Hall Institute, Melbourne, Australia); Melanie Price (University of Sydney, Sydney, Australia); Melissa Southey (University of Melbourne, Melbourne, Australia); Roger Milne (Cancer Council Victoria, Melbourne, Australia); Sid Deb (Melbourne Health, Melbourne, Australia); David Bowtell (Garvan Institute of Medical Research, Sydney, Australia).
